# Network models incorporating chloride dynamics predict optimal strategies for terminating status epilepticus

**DOI:** 10.1101/2024.08.13.607480

**Authors:** Christopher B. Currin, Richard J. Burman, Tommaso Fedele, Georgia Ramantani, Richard E. Rosch, Henning Sprekeler, Joseph V. Raimondo

**Author notes:** **Corresponding authors:** (CBC), (JVR).

## Abstract

Seizures that continue for beyond five minutes are classified as status epilepticus (SE) and constitute a medical emergency. Benzodiazepines, the current first-line treatment, attempt to terminate SE by increasing the conductance of chloride-permeable type-A GABA receptors (GABA_A_Rs). Despite their widespread use, benzodiazepines are ineffective in over a third of cases. Previous research in animal models has demonstrated that changes in intraneuronal chloride homeostasis and GABA_A_R physiology may underlie the development of benzodiazepine resistance in SE. However, there remains a need to understand the effect of these changes at a network level to improve translation into the clinical domain. Therefore, informed by data from human EEG recordings of SE and experimental brain slice recordings, we used a large spiking neural network model that incorporates chloride dynamics to investigate and address the phenomenon of benzodiazepine resistance in SE. We found that the GABA_A_R reversal potential (E_GABA_) sets SE-like bursting and determines the response to GABA_A_R conductance modulation, with benzodiazepines being anti-seizure at low E_GABA_ and ineffective or pro-seizure at high E_GABA_. The SE-like activity and E_GABA_ depended on a non-linear relationship between the strength of Cl^-^ extrusion and GABA_A_R conductance, but not on the initial E_GABA_ of neurons. Independently controlling Cl^-^ extrusion in the pyramidal and interneuronal cell populations revealed the critical role of pyramidal cell Cl^-^ extrusion in determining the severity of SE activity and the response to simulated benzodiazepine application. Finally, we demonstrate the model’s utility for considering improved therapeutic approaches for terminating SE in the clinic.

## INTRODUCTION

Most seizures terminate within a few seconds to minutes and do so spontaneously without the need for medical intervention. There are, however, some cases where seizure activity persists and when this lasts for more than 5 mins it is termed status epilepticus (SE) *(1)*. SE represents a medical emergency and if seizure cessation cannot be achieved is associated with significant morbidity and even mortality *(2)*. Current first-line treatment for SE recommends the use of benzodiazepines *(3)*. Benzodiazepines work by increasing the conductance of chloride (Cl^-^) permeable *γ*-aminobutyric acid (GABA) type A receptors (GABA_A_Rs), which mediate the majority of fast synaptic inhibition in the brain. The goal is to enhance inhibitory signalling to try to stop SE. Unfortunately, benzodiazepine therapy fails to halt seizures in over a third of patients, both adult and paediatric *(4–6)*, underscoring the critical need for a deeper understanding of the mechanisms behind benzodiazepine resistance in order to develop improved treatment strategies *(7)*.

Seizures reflect excessive excitation and synchronisation within the brain. Interneuronal populations, which release GABA and activate GABA_A_Rs on their synaptic targets, are a principal mediator of inhibition, which typically acts to prevent the initiation or spread of seizures *(8)*. The effect of fast GABAergic synaptic inhibition is dependent both on the magnitude of evoked GABA_A_R conductances (g_GABA_) and the underlying reversal potential for GABA_A_Rs (E_GABA_) *(9)*. Together these parameters control current flow through GABA_A_Rs and consequent shifts in neuronal membrane potential and firing activity (Figure 1A). Blockade of GABA_A_Rs using pentylenetetrazole, penicillin, picrotoxin and bicuculline are classically used to induce seizures both *in vitro* and *in vivo (10–15)* demonstrating a strong pro-seizure effect of reducing g_GABA_. In contrast, benzodiazepines act by enhancing the conductance of GABA_A_Rs following accompanying GABA binding *(16)*, and under typical conditions reduce the likelihood of seizures, demonstrating a typical anti-seizure effect of increasing g_GABA_ *(17)*.

**Figure 1:**
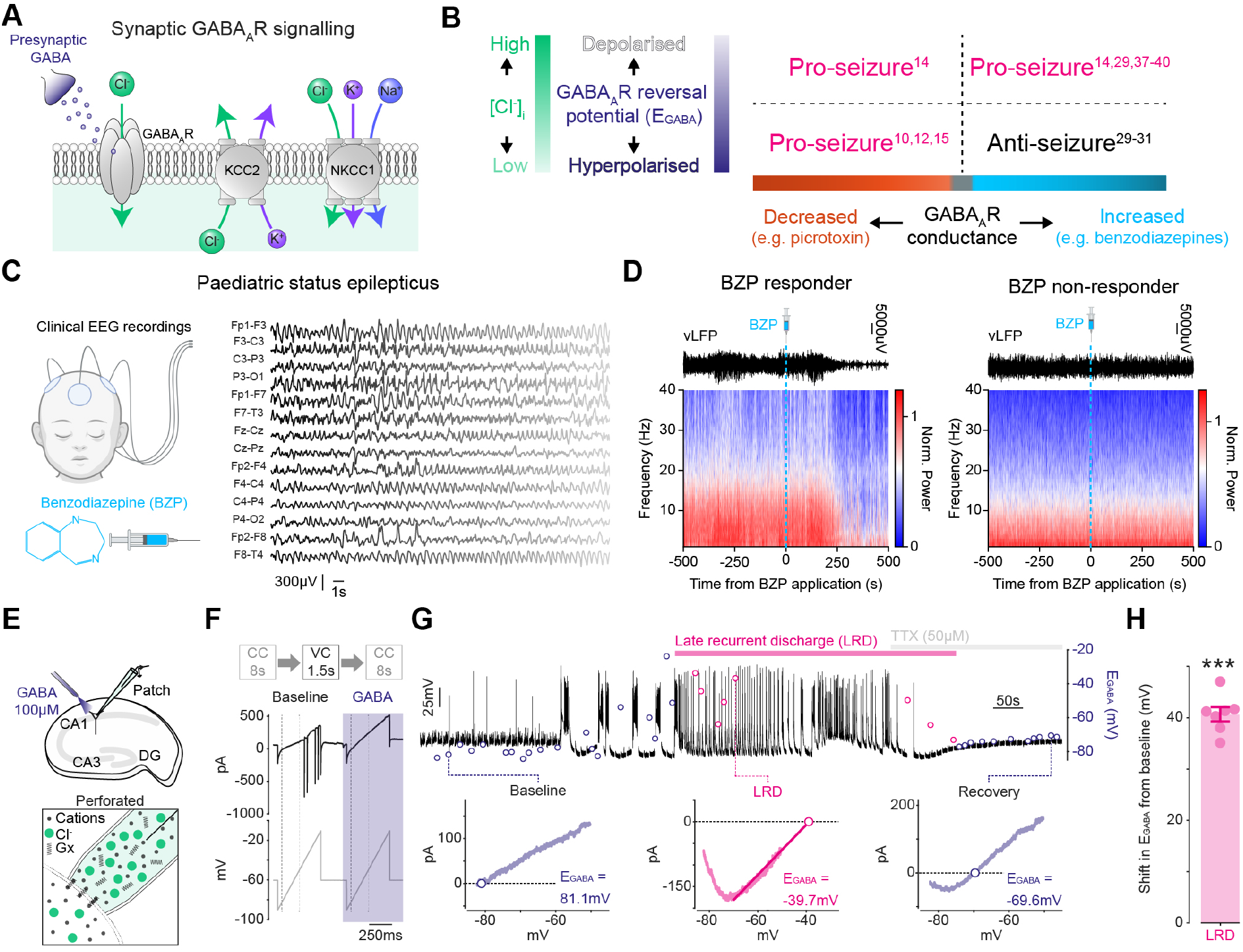
The GABA_A_R reversal potential determines the effect of GABA_A_R conductance manipulation on seizures. **(A)** Fast GABAergic synaptic inhibition is mediated by GABA_A_Rs, which are predominantly permeable to Cl^-^. Upon GABA binding, Cl^-^ flows down its electrochemical gradient depending on the reversal potential for GABA_A_Rs (E_GABA_) and the membrane potential. E_GABA_ is predominantly a function of the Cl^-^ gradient, which is modulated by the action of the Cl^-^ transporters KCC2 and NKCC1. **(B)** A table with references to experimental papers demonstrating a variable effect of GABA_A_R modulation on seizures depending on E_GABA_ *(10, 12, 14, 15, 29–31, 31, 37–40)*. Increasing GABA_A_R conductance with benzodiazepines can either be pro-seizure or anti-seizure depending on E_GABA_. **(C)** Clinical EEG recordings from a paediatric patient in status epilepticus *(data from 36***). (D)** Virtual local field potential recordings (vLFP) extracted from 21-channel EEG recordings with corresponding power spectra. Epochs consist of 500s on either side of benzodiazepine administration. One example shows a clear reduction in electrical activity following benzodiazepine administration (‘BZP responder’) whilst the other shows an example of a patient in which the benzodiazepine did not modify the EEG signal (‘BZP non-responder’). **(E)** Schematic depicting experimental setup of gramicidin perforated patch-clamp recordings from pyramidal cells in organotypic hippocampal brain slice cultures and accompanying somatic GABA application. **(F)** To measure E_GABA_ during seizure-like activity, the recording mode was rapidly switched from current-clamp (CC, 8 s duration) to brief periods in voltage clamp (VC, 1.5 s duration) every 10 s *(data from 14)*. While in VC, two consecutive voltage ramps (bottom trace) were applied: the first without GABA application and the second with GABA application (purple) directed toward the soma. The current was recorded (top trace) and a subtraction performed to calculate the GABA current and E_GABA_. **(G)** A representative recording from a CA1 pyramidal neuron where E_GABA_. measurements (dots) were made during the evolution of epileptiform activity in the 0 Mg^2+^ model (pink bar denotes the Late Recurrent Discharge phase, LRD, akin to Status Epilepticus). Dotted lines highlight different periods during the progression of epileptiform activity: baseline, during LRD, and following termination of activity / post-LRD with TTX (50 mM). Bottom, I-V plots were used to calculate E_GABA_. defined as the voltage at which the GABA current equals 0. **(H)** Population data demonstrating a profound shift in E_GABA_. from baseline (mean shift: 40.67 ± SEM 1.38 mV, N = 7, ***P<0.001, one-sample t-test).

As GABA_A_Rs are primarily permeable to Cl^-^, the transmembrane gradient for Cl^-^ sets E_GABA_ *(18)*. It is now well accepted that the intracellular concentration of Cl^-^, and hence E_GABA_, can change over multiple timescales as a function of the cumulative Cl^-^ fluxes through Cl^-^ transporters and channels *(19, 20)*. Cl^-^ transporters including the cation-chloride cotransporters NKCC1 and KCC2 utilise cation gradients to import and extrude Cl^-^ respectively, shifting the Cl^-^ gradient beyond a passive distribution *(20)*. Long-term changes in Cl^-^ cotransporter expression and function modifies steady-state E_GABA_ over development and in multiple disease states including epilepsy *(21, 22)*. In addition to these long-term changes, short-term changes in E_GABA_ can occur when Cl^-^ channels such as GABA_A_Rs are intensely activated causing Cl^-^ influx that overwhelms Cl^-^ extrusion mechanisms *(19). In vitro* and *in vivo* data from animal models has shown that this occurs during seizures and SE where intracellular Cl^-^ accumulation and a depolarising shift in E_GABA_ can reduce the inhibitory effectiveness of GABAergic interneuronal cell populations, or even render them excitatory *(23–25)*.

The modulation of g_GABA_ is commonly used to control seizure activity both in the clinic and the laboratory. Various data from patients and animal models have demonstrated that the effect of g_GABA_ modulation on seizures depends on the underlying transmembrane Cl^-^ gradient and E_GABA_. A reduction in g_GABA_, including via blockade of GABA_A_R using picrotoxin or bicuculline, is typically pro-seizure causing hyperexcitability regardless of E_GABA_ *(14, 26–28)* (Figure 1B). In contrast, enhancing g_GABA_ with positive allosteric modulators of GABA_A_Rs, such as benzodiazepines, can have an anti-seizure effect when intracellular Cl^-^ concentration and E_GABA_ are low *(29–31)* but can have no effect, or a pro-seizure effect when intracellular Cl^-^ concentration and E_GABA_ are high *(14, 29)* (Figure 1B). SE and how it is affected by pharmacological perturbation is the result of multiple dynamically interacting mechanisms between different cell-types in brain networks, which can be difficult to predict or to study experimentally. Computational models allow for simulations of the effects of individual parameter on neuronal dynamics, and therefore are an ideal tool to complement experiments for ascertaining the mechanistic underpinnings of the clinically relevant phenomenon of benzodiazepine resistant SE, and for designing improved therapeutic strategies. Previous computational models incorporating Cl^-^ dynamics have been successfully used to demonstrate the importance of Cl^-^ in affecting synaptic integration and information processing by single cells *(32–35)*.

Here we present the first large spiking neural network model that incorporates Cl^-^ dynamics. We use this model, informed by data from human electroencephalography (EEG) recordings of SE and experimental brain slice recordings, to better understand and address the phenomenon of benzodiazepine resistance in SE. We find that E_GABA_ establishes SE-like bursting in a network model and determines the response to GABA_A_R conductance modulation as observed experimentally with GABA_A_R antagonists and benzodiazepine. In addition, we determine that the network’s steady-state bursting activity and E_GABA_ depends on a non-linear interaction between the strength of Cl^-^ extrusion and GABA_A_R conductance, but not on the initial E_GABA_ of the neurons. Separately manipulating Cl^-^ extrusion in the pyramidal cell as opposed to interneuronal cell populations reveals the dominant role of pyramidal cell Cl^-^ extrusion in determining the intensity of SE-like activity and the response to simulated benzodiazepine application. Finally, we demonstrate the model’s utility for conceptualising therapeutic protocols for more rapidly terminating SE in the clinic.

## RESULTS

### The GABA_A_R reversal potential determines the effect of GABA_A_R conductance manipulation on seizures

To illustrate the clinical presentation of benzodiazepine-resistant SE, we extracted example EEG recordings from paediatric patients *(data from 36)*. A unique feature of these recordings is that they capture the pre-and post-effect of benzodiazepine application during SE. Notably in one patient, enhancing g_GABA_ with a benzodiazepine resulted in the cessation of the EEG readout of seizure activity over the course of minutes (‘BZP responder’, Figure 1C and D). However, in another patient, benzodiazepine application had no effect on seizure activity (‘BZP non-responder’, Figure 1D). While it is not currently feasible to measure intracellular Cl^-^ concentration or E_GABA_ in human patients, one can use animal models to study SE and use them as a proxy to gain mechanistic insights into how GABA_A_R physiology changes during persistent seizure activity. Here, we demonstrate from previous experimental data *(14)* that withdrawing Mg^2+^ from the perfusing solution of organotypic brain slice cultures can reproduce SE-like activity. Using this *in vitro* model, gramicidin perforated patch-clamp recordings are used to measure the E_GABA_ throughout the evolution of SE-like activity without perturbing intracellular Cl^-^ (Figure 1E and F). Through this data, we can observe how the E_GABA_ undergoes a significant depolarising shift when it enters a period of late recurrent discharges (LRD) that is electrographically similar to SE (Figure 1G and H). Stopping SE-like activity using tetrodotoxin returned E_GABA_ to more negative values. This seizure-associated shift in E_GABA_ explains why if a benzodiazepine is applied before or at the onset of a seizure in this model, the seizure-like activity can be prevented, delayed, or reduced *(14)*. However, if benzodiazepines are applied during status epilepticus-like activity, when E_GABA_ is elevated, network activity may remain unaffected or exacerbated *(14)*.

### A network model of status epilepticus is suppressed or enhanced by increased GABA_A_R conductance depending on the neuronal GABA_A_R reversal

To investigate the effect of E_GABA_ on how modulation of g_GABA_ affects seizure activity, we built a spiking neural network model consisting of leaky integrate-and-fire point neurons; 800 pyramidal cells (PC) and 200 interneurons (IN). These were interconnected and received a low level of constant, external excitatory drive (Figure 2A and Methods). E_GABA_ was set at the same constant value in all cell types. Incrementing E_GABA_ by 2 mV every 60 seconds of simulation from -74 mV until -42 mV showed that E_GABA_ strongly controls firing rate bursts in the network, with no bursting being observed with E_GABA_ less than -60 mV (Figure 2B and C). E_GABA_ values above -60 mV resulted in bursting comparable to the network bursts observed in experimental models of SE *(14)* and Figure 1G.

**Figure 2:**
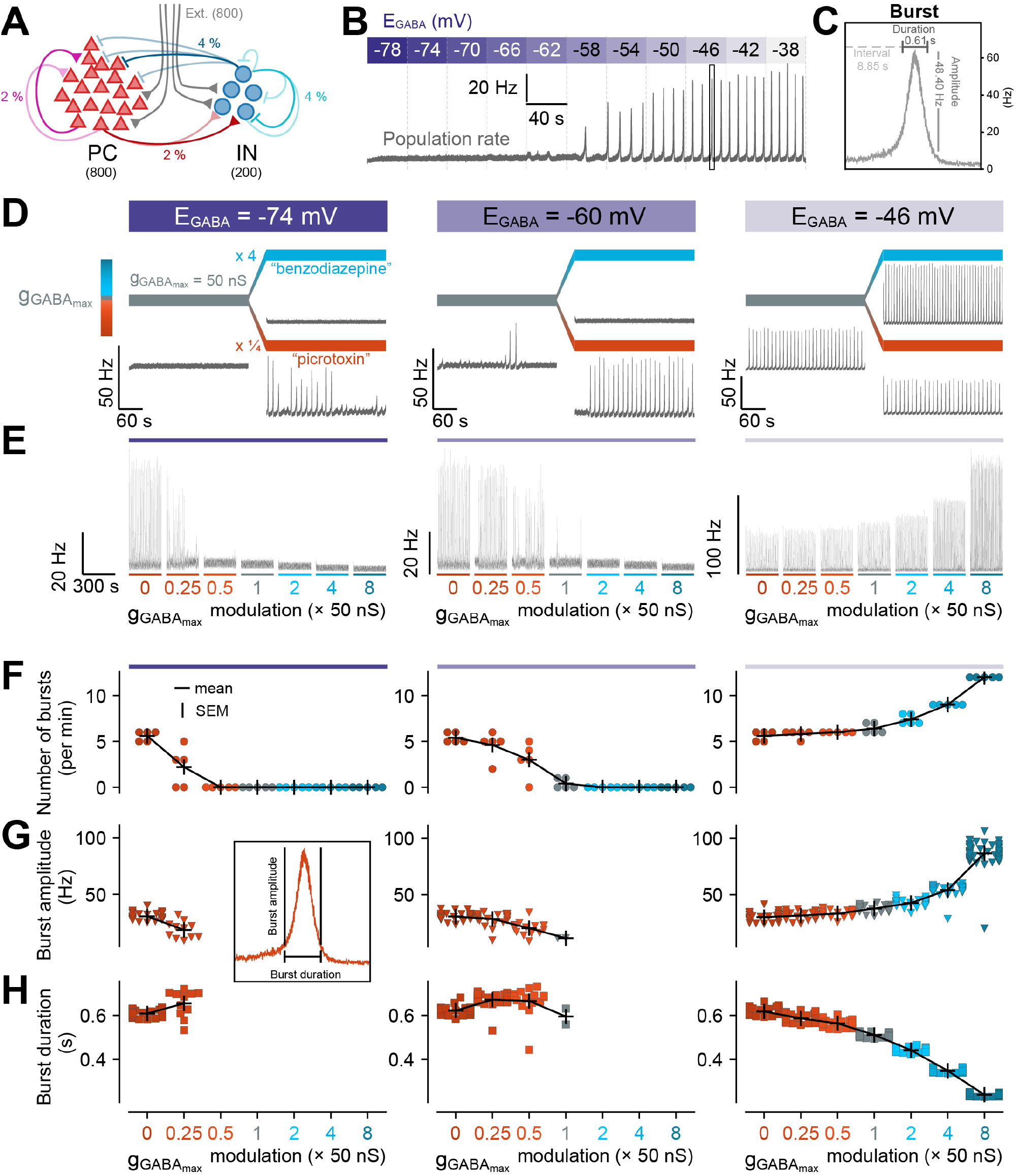
GABA_A_R reversal sets epileptiform bursting in a network model and determines the response to GABA_A_R conductance modulation. **(A)** Schematic of the spiking neural network model consisting of 800 pyramidal cells (PC) and 200 interneurons (IN), with 800 low frequency (2 Hz) and low weight (2 nS g_AMPAmax_) external inputs. The connection probabilities (in %) between and within populations are as follows: PC → PC: 2 %, PC → IN: 2 %, IN → IN: 4 %, IN → PC: 4 %. **(B)** E_GABA_ (purple scale bar) was altered at discrete time points, every 40 seconds, by 4 mV, from -78 mV until -38 mV. The population firing rate was monitored for bursts **(C)**, detected as twice the standard deviation of the mean, above 20 Hz and for at least 20 ms. **(D)** Average firing rate of neurons with hyperpolarising E_GABA_ (left: -74 mV), shunting E_GABA_ (middle: -60 mV) or depolarising E_GABA_ (right: -46 mV). After 300 s of normal GABA_A_R conductance (g_GABAmax_ = 50 nS, grey bar), g_GABAmax_ was either negatively (× ¼, simulating picrotoxin, orange bar) or positively modulated (× 4, simulating benzodiazepine, blue bar). Note simulated benzodiazepine application silencing bursting at a shunting E_GABA_ (middle), but exacerbating bursting when E_GABA_ is depolarised (right). **(E)** Population firing rates for a range of g_GABAmax_ modulations (as a proportion of 50 nS). **(F)** The number of bursts for each g_GABAmax_ modulation. Each coloured marker indicates the number of bursts per minute. Black crosses indicate the mean values. **(G)** The amplitude of oscillations for each g_GABAmax_ modulation. Each coloured marker indicates a burst. **(H)** The duration of bursts for each g_GABAmax_ modulation. Each coloured marker indicates a burst. For each simulation, E_GABA_ was kept constant over the entire 600 s duration, and g_GABAmax_ was modulated at 300 s. The population rate statistics were calculated over 300 s of the respective simulation of that condition.

Next, we sought to simulate the experiments described in Figure 1 by using our spiking neural network model to computationally determine how different neuronal E_GABA_ values might modify the effect of g_GABA_ modulation on seizure-like activity (bursts of increased population firing rate). To do so, after running the network simulation for 300 s with a “normal” g_GABA_ (g_GABAmax_) of 50 nS, g_GABAmax_ was altered to be either 4x smaller (e.g. modelling “picrotoxin” application, a GABA_A_R antagonist) or 4x larger (e.g. modelling “benzodiazepine” application, a positive GABA_A_R conductance modulator) (Figure 2D-H). To model the effects of different underlying E_GABA_ on this manipulation, the simulations were repeated using E_GABA_ of -74 mV (hyperpolarising), -60 mV (shunting) or -46 mV (depolarising). For hyperpolarising E_GABA_ (−74 mV, Figure 2D, left column), the network transitioned to occasional bursting following simulated picrotoxin application (12.5 nS g_GABAmax_) or remained quiescent following simulated benzodiazepine application (200 nS g_GABAmax_). At a shunting E_GABA_ (−60 mV, Figure 2D, middle column), the network transitioned from sporadic bursts to either continuous bursting following “picrotoxin” or was silenced following “benzodiazepine” application. Finally, at depolarising E_GABA_ (−46 mV, Figure 2D, right column), the network exhibited continuous bursting at baseline. The simulated application of picrotoxin by reducing g_GABAmax_ did not substantially change the network behaviour. However, positively modulating g_GABAmax_ (simulating application of a benzodiazepine) not only did not reduce bursting, but instead substantially increased it.

To examine the graded effect of g_GABAmax_ modulation, the procedure was repeated for a range of g_GABAmax_ values from 12.5 nS to 200 nS (Figure 2E). The number of bursts per min (Figure 2F), the amplitude of bursts (Figure 2G) and the duration of bursts (Figure 2H) were calculated. Together these results corroborate experiments results by demonstrating that at hyperpolarised E_GABA_s and shunting E_GABA_s, reducing g_GABA_ in the network elicits SE-like activity in the form of repeated bursting with increased g_GABA_ silencing the bursting activity. In contrast at a depolarised E_GABA_, increasing g_GABA_ increased the amplitude and frequency of bursting representing an exacerbation of SE-like activity.

### Chloride extrusion controls network bursting and the response to GABA conductance modulation

In the previous simulations intracellular Cl^-^ concentration, E_Cl_ and hence E_GABA_ were treated as static parameters. However, it is more accurate to consider these parameters as dynamic variables because intracellular Cl^-^ fluctuates as a function of activity-dependent Cl^-^ flux through GABA_A_Rs and Cl^-^ extrusion via the cation-chloride cotransporters such as KCC2 (Figure 3A). Therefore, in this set of simulations the evolution of E_GABA_ over time was modelled as a dynamic variable (see Materials and Methods). The efficacy of Cl^-^ extrusion in each neuron could be set by changing τ_KCC2_, the time constant of E_Cl_ (and hence E_GABA_) recovery, with slower τ_KCC2_ of 60 s or more representing reduced Cl^-^ extrusion by KCC2. We found that τ_KCC2_ determined the number of bursts in the network (Figure 3B and C) as well as the ultimate steady-state E_GABA_ (Figure 3B and D). Slower τ_KCC2_ values with resultant reduced Cl^-^ extrusion led to the network generating multiple bursts per minute together with elevated average steady-state E_GABA_. Whilst E_GABA_ was dynamic during simulations, changing from its initial value as well as rising and falling in response to individual network bursts (Figure 3B, inset), we found that the initial E_GABA_ (E_GABA0_) did not affect the final steady state E_GABA_ (Figure 3E and F), which was instead affected by the strength of Cl^-^ extrusion (τ_KCC2_).

**Figure 3:**
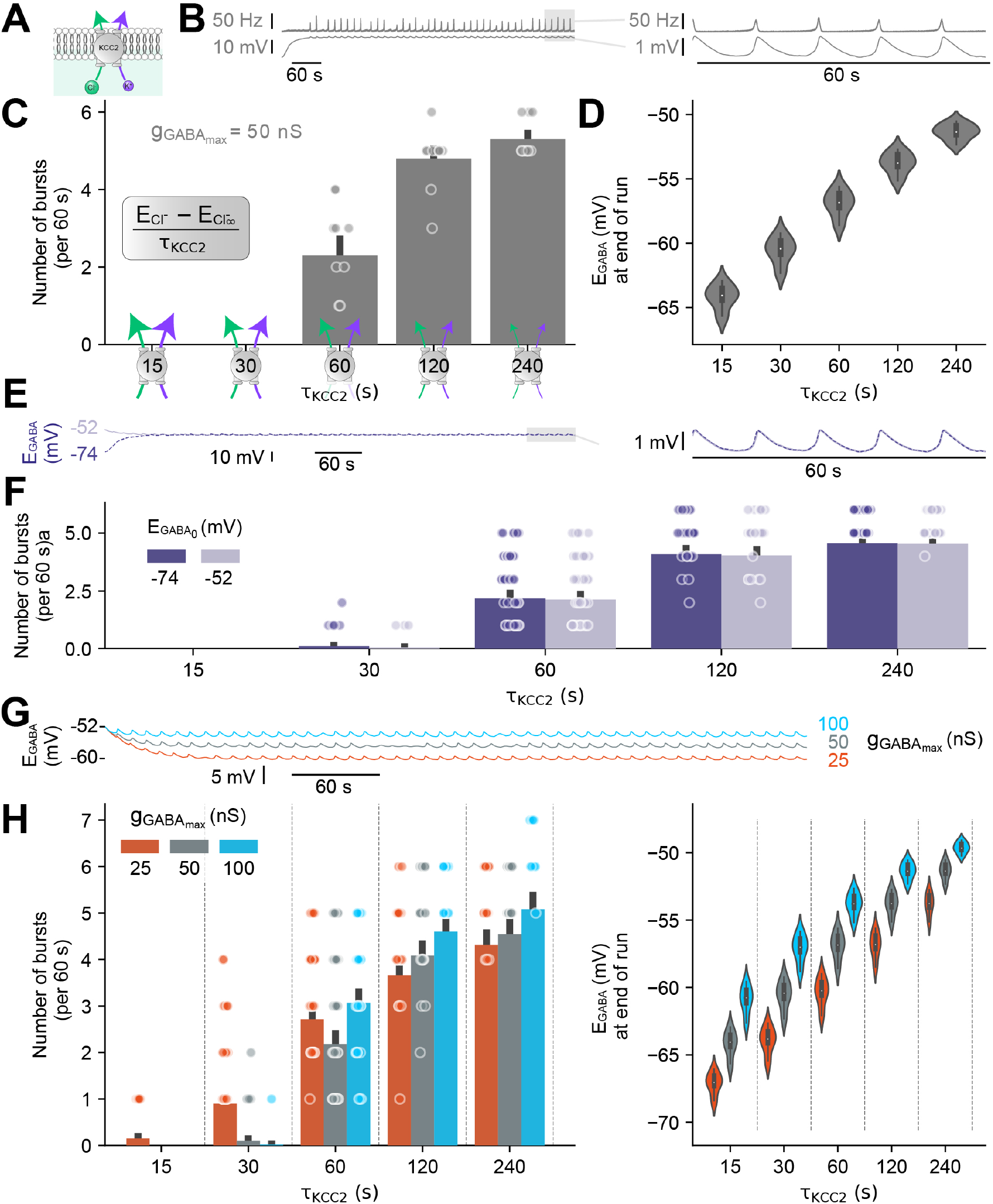
Chloride extrusion controls network bursting and the response to GABA_A_R conductance modulation. **(A)** Schematic of the primary Cl^-^ extrusion mechanism in adult neurons (KCC2) which is modelled as a single exponential decay to baseline (−88 mV) that depends on the time constant (τ_KCC2_). Smaller values of τ_KCC2_ indicate faster extrusion rates. **(B)** A simulation with dynamic Cl^-^ whereby E_GABA_ (bottom trace) depends on τ_KCC2_ (60 s) and the population activity (top trace) responds to elevated E_GABA_ by bursting. E_GABA_ initialised at -74 mV. Inset, zoom of traces showing ∼ 1 mV change in E_GABA_ in response to bursts. **(C)** The number of network bursts (per min) depended on the Cl^-^ extrusion strength (τ_KCC2_), with slower extrusion causing more bursts. **(D)** Reduced Cl^-^ extrusion (slower τ_KCC2_) resulted in elevated steady-state E_GABA_. **(E)** E_GABA_ did not depend on the initial E_GABA_ (E_GABA0_). For simulations with E_GABA0_ of either -74 mV (dark purple, dashed) or -51.6 mV (light purple, solid), the resulting steady-state E_GABA_ was the same. Inset, the traces for -74 mV and -51.6 mV E_GABA0_ are overlapping. Note that the same seed was used in both traces. **(F)** The number of bursts per min were independent of E_GABA0_. Black bars show the standard error of the mean (SEM). **(G)** E_GABA_ traces for different values of g_GABAmax_ (25 nS: orange, 50 nS: grey, 100 nS: light blue). All simulations started at -51.6 mV. **(H)** Histogram of the number of bursts per min and **(I)** violin plots of steady-state E_GABA_ for different values of g_GABAmax_ and τ_KCC2_. Each coloured marker indicates the number of bursts over the last 5 minutes of a 10-minute simulation (N = 10 simulations per condition). Violin plots include a box-and-whisker plot inside with the median value indicated as a white marker.

As Cl^-^ influx via activated GABA_A_Rs also affects E_GABA_ (see Figure 3B inset) we next sought to determine how modulation of g_GABAmax_ (akin to blockade or enhancement of GABA_A_Rs using picrotoxin or benzodiazepines respectively), might affect seizure-like activity in the context of dynamic Cl^-^ and E_GABA_. We first monitored E_GABA_ in networks with the same level of neuronal Cl^-^ extrusion (τ_KCC2_) but with different g_GABAmax_values of 25 nS, 50 nS, and 100 nS. Steady state neuronal E_GABA_ was substantially different between the conditions, with a difference of 9 mV between the low and high g_GABAmax_ conditions (Figure 3G). Next, to determine the interaction between Cl^-^ extrusion, GABA_A_R conductance and seizure-like activity, we systematically altered GABA_A_R conductance (g_GABAmax_) at different Cl^-^ extrusion rates (τ_KCC2_) and counted the number of network bursts (Figure 3H) together with measuring steady state E_GABA_ (Figure 3I). In networks with enhanced Cl^-^ extrusion (short τ_KCC2_) and resultant hyperpolarised steady state E_GABA_s, reduced GABA_A_R conductance (orange) increased bursting while enhanced GABA_A_R conductance (blue) silenced the networks. In contrast, in networks with low extrusion rates (long τ_KCC2_) and depolarised steady state E_GABA_s, reducing GABA_A_R conductance reduced bursting whilst increasing GABA_A_R conductance (akin to benzodiazepine application) exacerbated bursting.

### Compromised chloride extrusion in the pyramidal cell population is the major determinant of network bursting

Thus far, to determine the effects of compromised Cl^-^ extrusion on SE-like activity as represented by network bursting, we have manipulated Cl^-^ extrusion (by adjusting τ_KCC2_) in all neurons. To investigate how Cl^-^ extrusion in specific neuronal subpopulations might affect SE-like activity, we altered Cl^-^ extrusion either in the pyramidal cell (PC) or interneuronal (IN) populations alone. With a view to understanding potential cell-type specific modulation of Cl-extrusion to affect SE, in these simulations E_GABA_ was initially set at a depolarised, SE-like level of -51.6 mV and allowed to evolve dynamically thereafter. Using a “baseline” g_GABAmax_ of 50 ns, it was immediately apparent that Cl^-^ extrusion in pyramidal cells 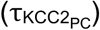 strongly determined bursting activity (Figure 4A, B and D). Strong Cl^-^ extrusion in pyramidal cells 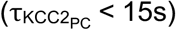 terminated network bursts whilst progressively weaker Cl^-^ extrusion resulted in increased bursting (Figure 4B, D). In comparison, modulation of Cl^-^ extrusion exclusively in the GABAergic interneuronal population 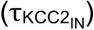 had a substantially smaller effect on bursting activity (Figure 4C and D).

**Figure 4:**
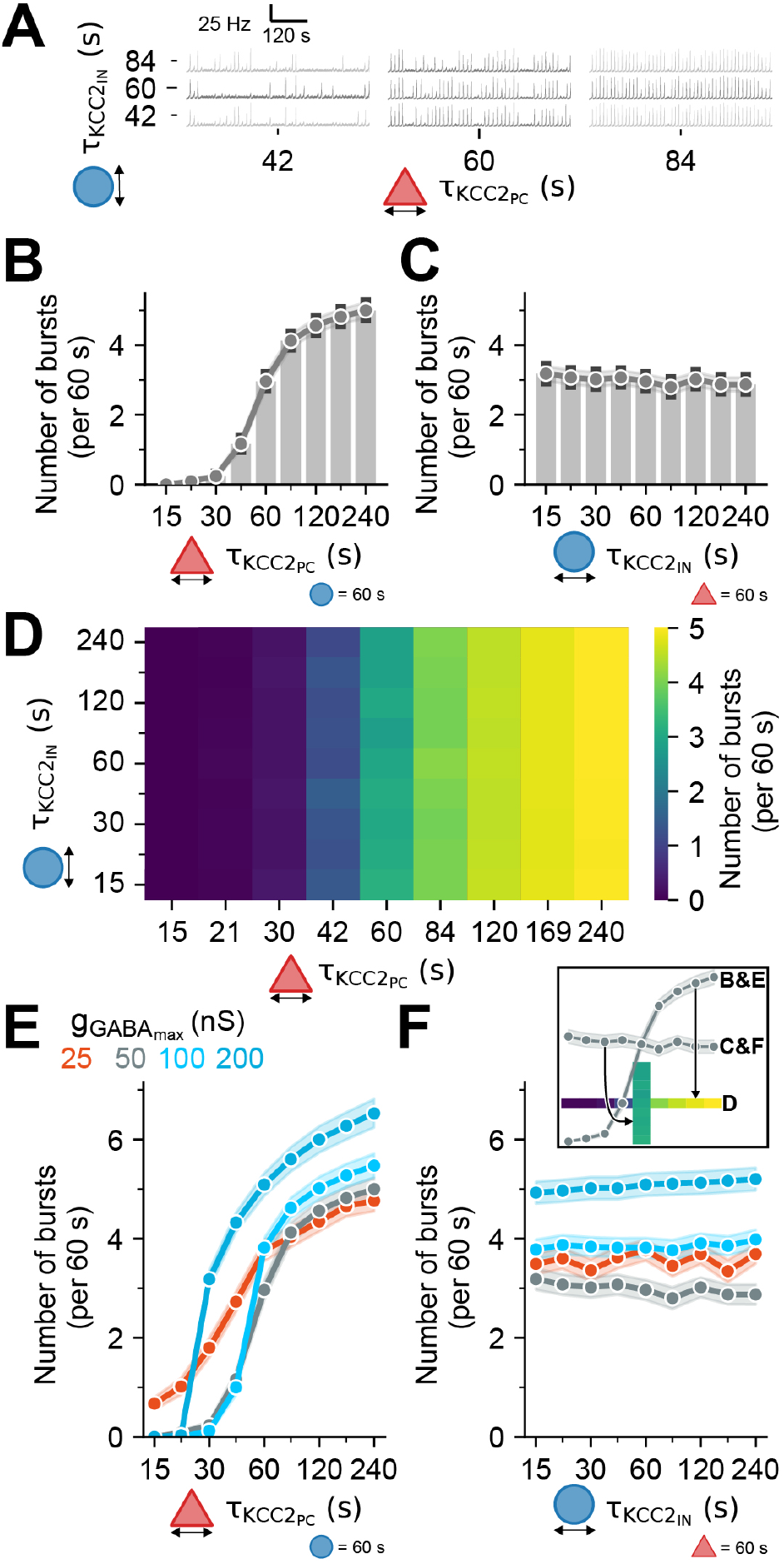
Compromised chloride extrusion in the pyramidal cell population is the major determinant of network bursting. **(A)** Population firing rate traces where τ_KCC2_ was independently varied for the pyramidal cell 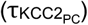 and interneuron 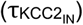 populations respectively. For all simulations, E_GABA_ was initialised at -51.6 mV. **(B)** Histograms of the number of bursts versus 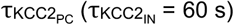 or **(C)** 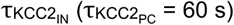. Black bars indicate ± SEM, N = 5 simulations. **(D)** Heatmap of the average number of bursts for a matrix of 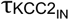 and 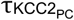 values. The average is calculated from 5 separate simulations. **(E)** Line plots of the number of bursts versus 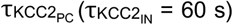 or 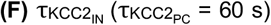, for values of 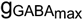 (25 nS: orange, 50 nS: gray, 100 nS: light blue, 200 nS: darker blue). Shaded areas indicate SEM, N = 5 simulations. The inset shows the relation between the heatmap and the bar and line plots.

The strength of GABA_A_R conductance modulated the effect of cell type specific Cl^-^ extrusion on network bursting (Figure E and F). Increasing g_GABAmax_ (Figure E) decreased the number of bursts at strong (i.e. short) 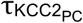 but increased the number of bursts substantially at weak (i.e. slow) 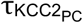. This was in line with the previous results. Although modulation of 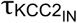 had much smaller effects on network bursting, trends could be observed particularly following manipulation of g_GABAmax_. At a baseline g_GABAmax_ of 50 nS (grey), the total number of bursts decreased with slower 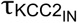. In contrast at a high g_GABAmax_ the number of bursts increased with slower 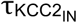 (Figure F). Overall, these results highlight the importance of the PC population’s Cl^-^ extrusion capacity, over that of the IN population, for managing network activity and preventing SE-like activity.

### Interacting chloride plasticity mechanisms determine the optimal approach for terminating network bursts

So far, we separately compared how SE-like activity in the form of network bursting is affected by GABA_A_R reversal potential (E_GABA_), GABA_A_R conductance (g_GABAmax_) and neuronal Cl^-^ extrusion (τ_KCC2_). Here, we investigated how all three of these variables interact in a way that could help guide future strategies for medical intervention in SE (Figure 5). First, we used a network with static Cl^-^ where E_GABA_ was held constant at different values. Positively modulating g_GABAmax_ from a baseline of 0 nS g_GABAmax_ decreased the number of bursts when E_GABA_ < -52 mV but increased the number of bursts when E_GABA_ > -52 mV (Figure 5A). This showed that g_GABAmax_ modulation had different trajectories for different E_GABA_s. If initial g_GABAmax_ modulation did not reduce the rate of bursting, further g_GABAmax_ modulation only served to exacerbate network activity further. This suggests that in the clinic, having EEG feedback to determine the response to initial benzodiazepine treatment could be useful for determining next treatment steps, particularly where seizure activity is not aborted.

**Figure 5:**
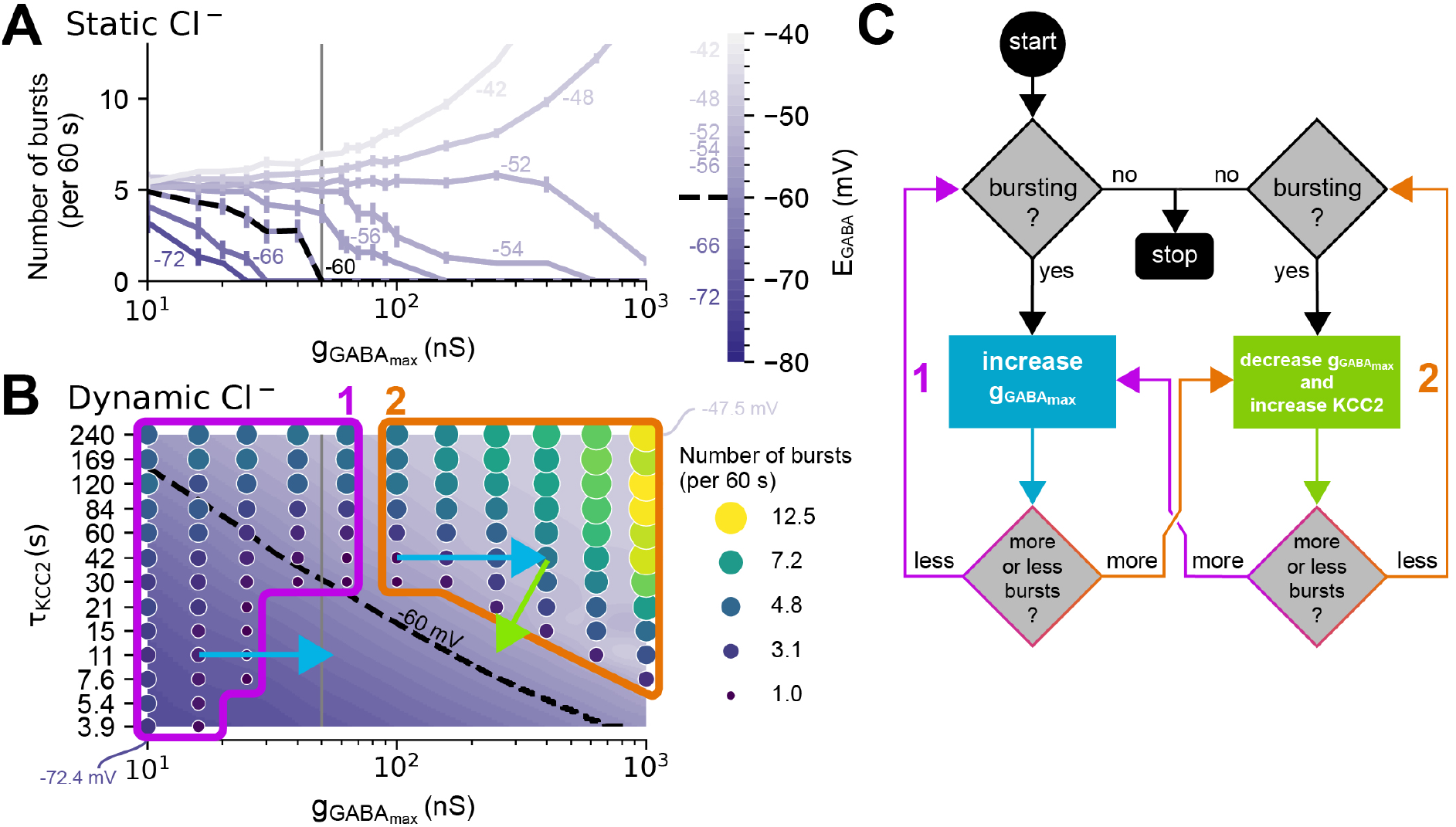
Interacting chloride plasticity mechanisms determine the optimal approach for terminating network bursts. **(A)** For simulations with static Cl^-^, the effect of positively modulating GABA conductance (g_GABAmax_) on a network’s bursting frequency depends on E_GABA_ (purple lines ± SEM, N = 10 simulations per condition, 60 s each). E_GABA_ was varied from -74 mV to -42 mV in increments of 2 mV. **(B)** Burst frequency as a function of g_GABAmax_ and τ_KCC2_ in simulations with dynamic Cl-, plotted together with average E_GABA_. E_GABA_ was measured at either the time of a burst or after 600 s if there were no bursts. Dashed line represents E_GABA_ = -60 mV, with the colourmap displayed in discrete 1 mV intervals. Low g_GABAmax_ elicits bursting behaviour regardless of τ_KCC2_. Increasing g_GABAmax_ decreases bursts if τ_KCC2_ is fast enough. Application of positive g_GABAmax_ modulation (blue arrows, simulating benzodiazepine application) decreases bursting at fast (short) τ_KCC2_ indicating that the network is in Regime 1 (magenta) but increases bursting at slow (long) τ_KCC2_, indicating that the network is in Regime 2 (orange). Recovery from bursting in Regime 2 can be facilitated by reducing g_GABAmax_ and increasing Cl^-^ extrusion (green arrow). **(C)** Decision tree depicting the optimal strategy for terminating seizures in SE based on the simulations in ‘B’.

To extend this idea further, we included the more realistic scenario of dynamic Cl^-^ and how neuronal Cl^-^ extrusion complicates the relationship between GABA_A_R conductance and synchronised bursting activity in the network. This is important because τ_KCC2_ and g_GABAmax_ together control to what extent bursting activity raises E_GABA_ in a positive feedback loop (see Figure 3H and the equation for 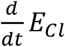 in Materials and Methods). By plotting the number of network bursts as a function of g_GABAmax_ and τ_KCC2_ together with E_GABA_, useful observations were made. Firstly, different combinations of τ_KCC2_ and g_GABAmax_ determine whether a network bursts and whether increasing g_GABAmax_ will decrease or increase bursting (Figure 5B). Given a network which is bursting (or a patient in SE), with no other information, it is not possible to know what the likely effect of increasing GABA_A_R conductance (i.e. benzodiazepine treatment) will be. However, by measuring the response (extent of network bursting) to increasing g_GABAmax_ (“benzodiazepine treatment”, blue arrows), one can then determine where in the network “landscape” one is positioned to choose the optimum next step to increase the likelihood of terminating seizures. We summarise this as a decision tree in Figure 5C. If raising g_GABAmax_ (blue arrow) reduces bursting, then the network is in Regime 1 (magenta). g_GABAmax_ can then be raised further. If raising g_GABAmax_ increases bursting, then the network is in Regime 2 (orange). In this case, g_GABAmax_ should be reduced by halting further benzodiazepine treatment. In addition, neuronal Cl^-^ extrusion should be increased if possible (green arrow). We note that pharmacological enhancers of Cl^-^ extrusion are not yet clinically available. Nonetheless, our modelling results demonstrate how in the case that GABA_A_R conductance increases network activity in SE, an optimal strategy for suppressing persistent seizures would be to reduce GABA_A_R conductance together with enhancing Cl^-^ extrusion.

## DISCUSSION

In this study we used novel computational models informed by clinical and experimental data to investigate the phenomenon of benzodiazepine resistance in status epilepticus (SE). Here we have implemented observations from experiments in a large spiking neural network model, and demonstrate the effects of chloride dynamics. Given recent experimental demonstrations of profound Cl^-^ fluctuations during seizures and SE *(14, 41)*, this is a necessary next step toward understanding benzodiazepine resistance in SE. Our findings align with experimental evidence, showing that the impact of GABA_A_R conductance (g_GABAmax_) modulation on SE-like activity is determined by neuronal GABA_A_R reversal potential (E_GABA_). Further, we demonstrate that compromised Cl^-^ extrusion within pyramidal cells is a major determinant of network bursting, and that a considered modulation of both GABA_A_R conductance and Cl^-^ extrusion is optimal for arresting SE-like activity.

We constructed our computational spiking neural network model to align with the well-characterised *in vitro* 0 Mg^2+^ brain slice model of acute, convulsive SE. As during *in vitro* experiments, transitioning the network to recurrent firing-rate bursts representing SE-like activity is linked with an elevation of E_GABA_ (i.e. raised [Cl^-^]_i_). Network bursting is affected by multiple factors including the properties of neurotransmitter release, receptor conductances, network size and connectivity *(42, 43)* (see also S3). Nonetheless, we could confirm that E_GABA_ was always a principal factor in transitioning a network from stable to bursting, SE-like behaviour.

The application of *in silico* pharmacological treatment to modulate g_GABAmax_ corroborated experimental results and highlighted the contrasting behaviour of increasing g_GABAmax_, which is anti-seizure at low, physiological E_GABA_, but pro-seizure at high, pathological E_GABA_ *(44, 45)*. In our model, the “edge” for transitioning between these two respective effects was an E_GABA_ of -60 mV. Experimental data suggest that the longer seizure activity continues unabated, the more likely it is that neurons undergo intracellular Cl^-^ accumulation and a positive shift in E_GABA_ *(14, 46, 47)*. This, together with our modelling data, helps provide a mechanistic explanation for the clinical observation that patients with SE who seize for longer prior to initial treatment are more likely to be resistant to benzodiazepine treatment *(7, 14, 48)*.

Previous work has shown that the continued seizure activity that occurs during SE results in internalisation of GABA_A_Rs *(49, 50)*, which both progressively exacerbates seizures and impairs the potential effectiveness of benzodiazepines as anti-seizure agents. This is because the ability of benzodiazepines to increase g_GABAmax_ is compromised by a lack of GABA_A_Rs with the requisite subunits *(51)*. Until recently this has been suggested as the major mechanism underlying benzodiazepine resistance in SE. Our model shows the importance of both changes in g_GABAmax_ together with activity-dependent dynamics in Cl^-^ and E_GABA_ for predicting the effectiveness of benzodiazepine resistance in SE.

A powerful determinant of how readily neuronal E_GABA_ increases following activity-dependent Cl^-^ influx through GABA_A_Rs is the strength of neuronal Cl^-^ extrusion via cotransporters such as KCC2. Increasing Cl^-^ extrusion can increase the seizure threshold, help terminate seizures or prevent them altogether *(45, 52, 53)* while blocking KCC2 can allow seizures to start spontaneously *(54)*. In agreement with this, in our simulations that accounted for Cl^-^ accumulation through GABA_A_Rs and Cl^-^ extrusion through KCC2, we found that simulating sufficiently slow Cl^-^ extrusion in all neurons could cause a network to start bursting without further external provocation. Further, at fast Cl^-^ extrusion rates which resulted in low steady-state E_GABA_, increasing g_GABAmax_ reduced bursting, while at slow Cl^-^ extrusion rates, which resulted in high steady state E_GABA_, enhancing g_GABAmax_ exacerbated bursting. Here, the steady-state E_GABA_ was not dependent on initial conditions. That is, the network’s steady-state E_GABA_ ultimately reached the same value regardless of whether it started at physiological (−74 mV) or pathological (−51.6 mV) E_GABA_. This result affirms the importance of Cl^-^ extrusion for affecting network excitability.

By selectively altering Cl^-^ extrusion in each population of neurons (pyramidal vs interneuronal cells), we determined that Cl^-^ extrusion in pyramidal cells is the predominant factor for controlling SE-like activity while Cl^-^ extrusion in interneurons had only a minor influence. The failure of inhibitory connections from interneurons to pyramidal cells plays a key role in causing persistent bursting. The network was in SE-like activity if Cl^-^ extrusion in pyramidal cells was slow enough, regardless of Cl^-^ extrusion in interneurons. This suggests that focusing on reducing pyramidal cell Cl^-^ accumulation is a major avenue through which seizure activity can be arrested. This has been shown experimentally where overexpression of KCC2 in cortical pyramidal cells reduced the seizure promoting effect of excitatory interneuronal signalling *(25)*.

While ours is the first large, spiking neural network to simulate Cl^-^ dynamics, and the first to apply this to model SE, previous computational models have explored the relevance of ion dynamics in SE using two-cell biophysical or mean-field approaches *(55–57)*. Our findings are in line with this work by reiterating the connection between raised intracellular Cl^-^ accumulation and E_GABA_ in maintaining extended seizures. Our model did not simulate dynamics in other ions including K^+^, Na^+^, H^+^ and 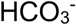, which are also known to both modulate and be modulated by seizure activity *(58)*. These ions indirectly interact with Cl^-^ through atleast two mechanisms: co-transporters like KCC2, which couple Cl^-^ transport to K^+^, and receptors such as GABA_A_Rs, which are permeable to both Cl^-^ and 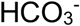. Incorporating these additional biophysical details could enhance the accuracy of our simulations. In the context of a large network model such as this, however, it would come at a potentially prohibitive computational cost. Modelling the dynamics of other ions could also identify or confirm other, Cl^-^ independent mechanisms for aborting seizures. For example, enhancing intraneuronal H^+^ (acidosis) is known to be anti-seizure via various mechanisms *(58)*. Further, raised intraneuronal Na^+^ has also been identified as contributing seizure termination *(56)*.

It is worth noting that our spiking neural network model was uniformly connected without explicitly modelling space or the propagation of activity through different brain regions. It did not capture potential spatial or inter-regional dynamics of activity-dependent shifts in E_GABA_. Differences in neuronal E_GABA_ between brain regions could explain why many people with SE still respond to first-line treatment with benzodiazepines *(7)*. In these patients it is possible that actively seizing networks with raised E_GABA_ are surrounded by less-affected areas with low E_GABA_ where benzodiazepines still enhance inhibition. The combined effect of a benzodiazepine likely depends on the extent to which various brain areas have been recruited into a seizure. Future work could extend our model by incorporating spatial considerations. This consideration does not affect our major conclusion here, namely that should SE not abate following the initial delivery of a high dose of benzodiazepines, serious consideration should be paid to not repeatedly delivering further agents of this class. Second line treatment that also engages other non-Cl^-^ linked inhibitory systems should rather be considered. As one of multiple potential examples, this could include phenobarbital, which in addition to its effect on GABA_A_Rs, is also an effective antagonist of AMPA and kainate glutamatergic receptors at higher concentrations *(59, 60)*.

In summary, our simulations provide an experimentally supported method of investigating mechanisms in SE. Our findings suggest that co-targeting of pyramidal cell extrusion with appropriate GABA_A_R conductance modulation represents a powerful strategy for terminating SE. Although several pharmacological agents which enhance Cl^-^ extrusion via increasing the activity of KCC2 have been identified *(53)*, none have been cleared for clinical use. Regardless, this research reveals a promising avenue for future advances in the management of benzodiazepine resistant SE.

## MATERIALS AND METHODS

### *In vivo* paediatric EEG

Details of clinical recordings are fully explained in previous work *(35)*. In short, anonymised patient data was acquired retrospectively and was approved by the local ethics committee (Kantonale Ethikkommission Zürich, KEK-ZH PB-2020-02580). We extracted data from paediatric patients (under 18 years) with SE who underwent scalp EEG recordings between July 2008 and February 2020 at the University Children’s Hospital Zurich. Clinical EEG recordings (21 electrodes, international 10–20 electrode layout, 256-Hz sampling) were reviewed and for the purposes of this study, illustrative examples of response to benzodiazepines were selected. To demonstrate the effect of benzodiazepine on the EEG signal, we isolated a 1000s epoch which included a 500s window on either side of the administration. A virtual local field potential (vLFP) was extracted using Statistical Parametric Mapping (SPM12) to show the overall effect of benzodiazepine on the patients EEG. Spectrograms were generated using Morlet wavelets from the PyWavelets library *(61)*.

### *In vitro* brain slice gramicidin perforated patch clamp

Details of *in vitro* brain slice recordings are fully explained in previous work *(14)*. Briefly, organotypic hippocampal slice cultures were prepared from mice. The use of animals was approved by the University of Cape Town Animal Ethics Committee. Recordings were performed 6–14 days post culture, equivalent to postnatal days 13 to 21 in mice and a paediatric age range in humans. Gramicidin perforated patch-clamp recordings were performed from CA1-CA3 hippocampal pyramidal cells using glass pipettes containing a high Cl^-^ internal solution containing (in mM): KCl (135), NaCl (8.9), HEPES (10) and 80 mg/ml gramicidin. The standard artificial CSF was composed of (in mM): NaCl (120); KCl (3); MgCl_2_ (2); CaCl_2_ (2); NaH_2_PO_4_ (1.2); NaHCO_3_ (23); D-glucose (11) with pH adjusted to be between 7.35 and 7.40 using 0.1 mM NaOH. To elicit SE-like activity, slices were perfused with aCSF lacking Mg^2+^ *(62, 63)*. To measure E_GABA_ during SE-like activity, voltage ramps in voltage-clamp mode with and without GABA (100 μM) application directed to the cell soma were interleaved between measurements of spontaneous activity recorded in current-clamp mode *(14, 64)*.

### *In silico* model of status epilepticus

We constructed a network of 1000 leaky integrate-and-fire point neurons with 20% as the GABAergic interneuron (IN) population and 80% as the glutamatergic pyramidal cell (PC) population *(65)*. Populations differed by their membrane capacitance, refractory period, and volume (see Table 1). The network had 2% connectivity from the PC population, including recurrent connections, and 4% connectivity from the IN population, also including recurrent connections, and 800 low frequency (2Hz) Poisson inputs for sustained activity *(43, 66)*.

**Table 1:**
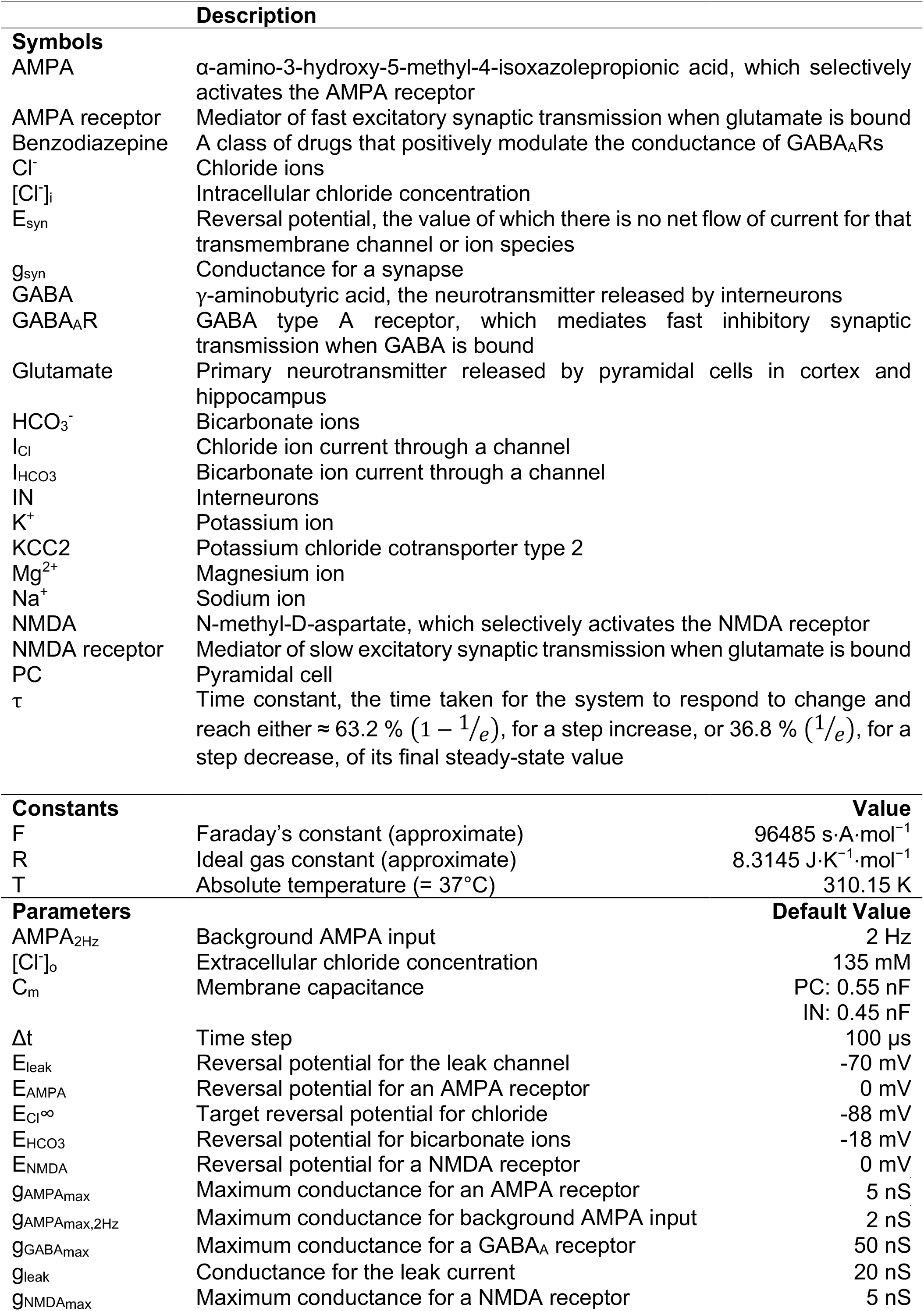
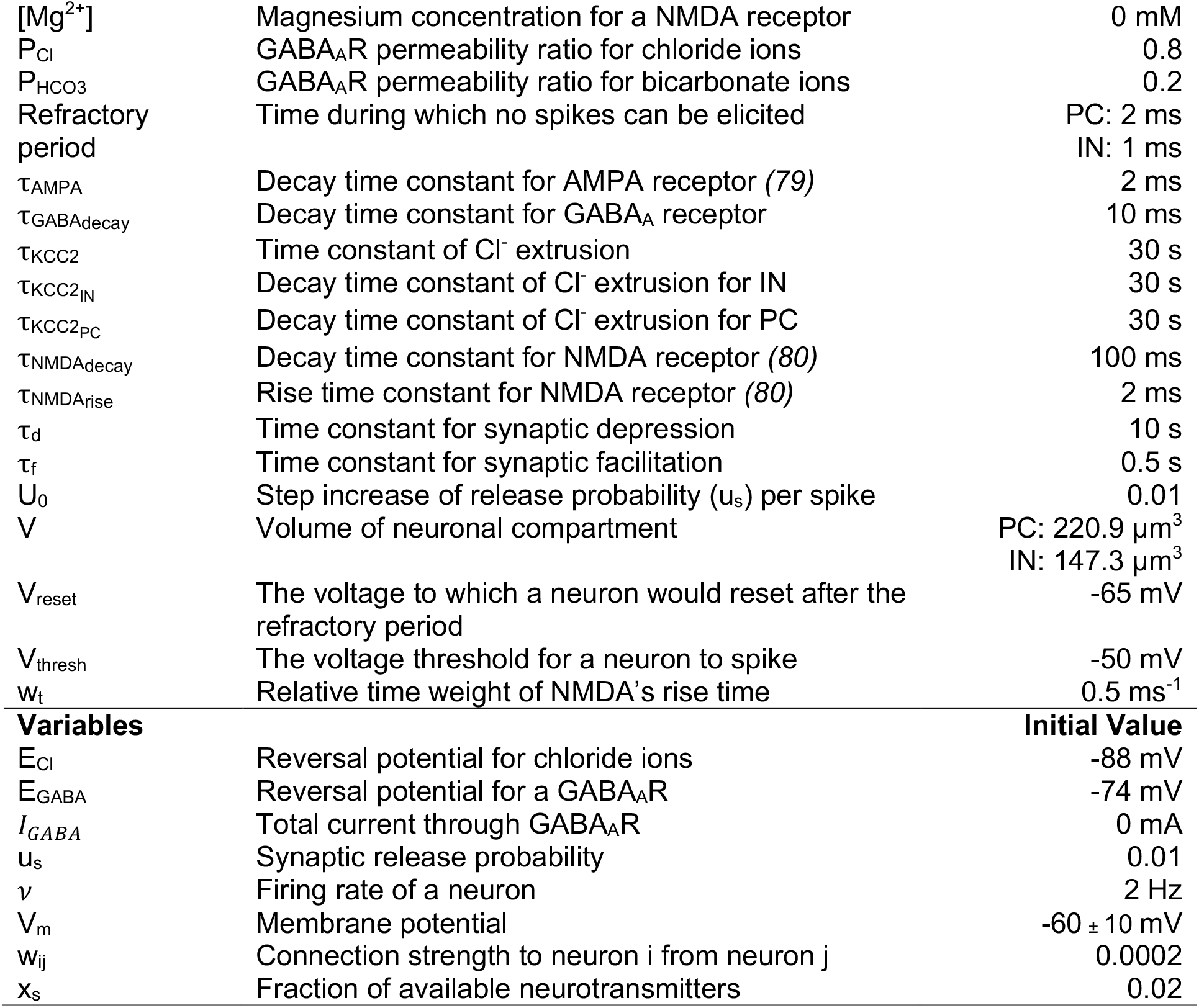
Symbols, Constants, and Parameters.

Change in the neuronal membrane potential (V_m_) was governed by

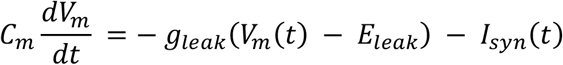

where g_leak_ (20 nS) and E_leak_ (−70 mV) are the conductance and reversal potential for the passive membrane channels respectively, C_m_ is the membrane capacitance (PC: 0.55 nF, IN: 0.45 nF) and I_syn_ is the synaptic current. Once threshold (−50 mV) was reached, a neuron would spike and reset (−65 mV) with a refractory period (PC: 2 ms, IN: 1 ms) during which a neuron could not spike again. Several mechanisms can enable a network to have bursts that occur over hundreds of milliseconds and seconds, including NMDA *(67–69)* and short term synaptic plasticity *(42, 70, 71)*, which were both incorporated.

Each neuron received synaptic input currents from AMPA, NMDA, and GABA_A_ receptors, as well as low frequency (2 Hz) background AMPA input (max conductance of 2 nS), modelled according to

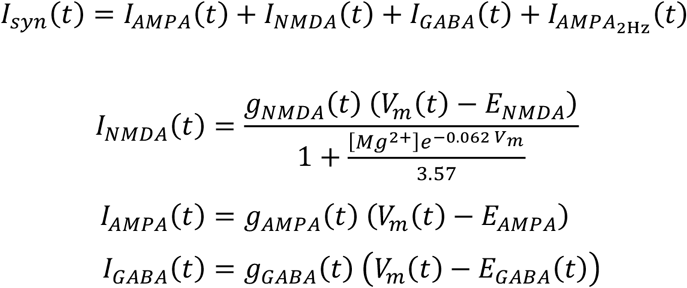

where E_syn_ is the reversal potential for that synapse, g_syn_ is the conductance for that synapse, and [Mg^2+^] is the magnesium block which was set to 0 mM within 5 seconds to mimic the *in vitro* experiments. The reversal potentials were E_NMDA_ = 0 mV, E_AMPA_ = 0 mV, and

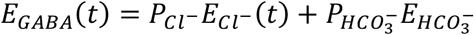

where E_Cl_ = -88 mV (for static Cl^-^ simulations), E_HCO3_ = -18 mV, and channel permeability ratios were P_Cl_ : P_HCO3_ = 0.8 : 0.2 such that E_GABA_ = -74 mV. The chord conductance formulation for the GABA_A_R reversal potential was used for its computational efficiency when modelling dynamic E_Cl_. For dynamic Cl^-^ simulations, E_Cl_ was modelled according to

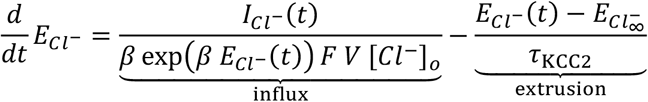

with

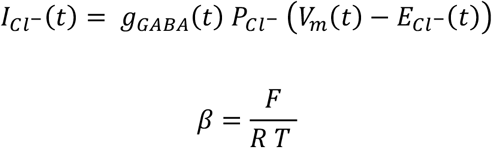

where R is the ideal gas constant, T is the temperature in Kelvin, F is Faraday’s constant, V is the neuronal volume (PC: 220.9 μm^3^, IN: 147.3 μm^3^), [Cl^-^]_o_ is the external Cl^-^ concentration, I_Cl_ is the Cl^-^ current through a GABA_A_R, E_Cl_∞ is the target reversal potential for Cl^-^ (−88 mV) and τ_KCC2_ is the time constant for KCC2. See Supplementary Information for derivation details of influx, and extrusion was modelled akin to *(72)*.

The change in synaptic conductance was modelled as a single-or double-exponential curve. AMPA and GABA_A_ synapses were modelled as

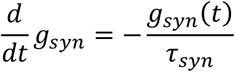

where τ_syn_ is the decay time constant (τ_AMPA_ = 2 ms, τ_GABA_ = 10 ms). NMDA conductance was modelled as

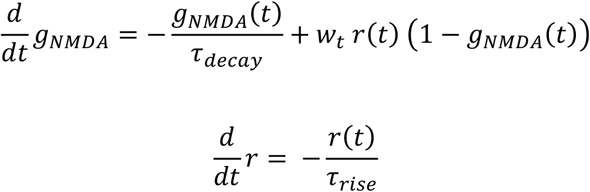

where τ_decay_ is the decay time constant for NMDA (100 ms), τ_rise_ is the rise time constant for NMDA (2 ms), r is the exponential rise component of the conductance, and w_t_ is its relative time weight (0.5 ms^-1^). Incoming synaptic conductances were calculated according to

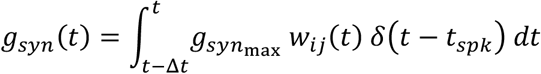

where 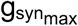 is the max synaptic conductance 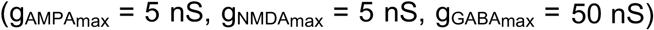, w_ij_ is the connection strength (‘weight’) for neuron i from neuron j (see details below), δ is the Dirac delta function, t_spk_ is the spike time of the presynaptic neuron, and Δt is the integration time window (100 μs).

Synaptic signalling involved short-term plasticity (STP) with depression – a depletion in the amount of neurotransmitter available (x_s_) – and facilitation – an increased release probability of neurotransmitters (u_s_). The connection strength of a synapse (w_ij_) was, therefore, defined according to

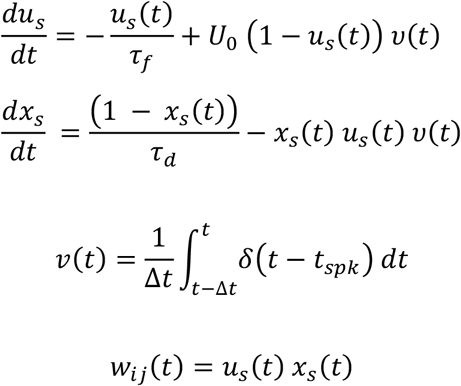

where U_0_ is the step increase in u_s_ per spike (0.01), τ_f_ is the time constant for facilitation (0.5 s), τ_d_ is the time constant for depression (10 s), and υ is the firing rate of presynaptic neuron j for a given time window, Δt (100 μs).

Simulations were carried out using the Brian2 Python package *(73)*, with C++ standalone code generation. Brian2’s default time step of 100 μs and the default numerical integration method of forward Euler were used. E_GABA_ was averaged over simulations (n >= 5) from at least one PC and IN per simulation. A burst was detected if the population firing rate was above 20 Hz and two standard deviations above baseline for at least 20 ms.

All model source code (available online at https://github.com/ChrisCurrin/goldilocks-GABA) was written in Python with data processing performed using NumPy *(74)*, SciPy *(75)*, and Pandas *(76)*. Results were plotted using the Matplotlib and Seaborn libraries *(77, 78)*.

## SUPPLEMENTARY INFORMATION

### S1. Efficiently modelling the change in chloride ion reversal potential due to ion influx via GABA_A_Rs

Instead of modelling the change in chloride ion (Cl^-^) reversal potential (E_Cl_) as dependent on the internal chloride ion concentration ([Cl^-^]_i_) due to Cl^-^ influx via GABA_A_R, it is more computationally efficient to calculate the time derivative of 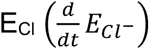 directly.

The Nernst equation for Cl^-^, moving the ‘*z* = −1’ into the logarithm, is

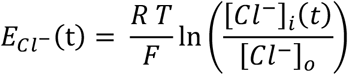

which we differentiate, while remembering that [Cl^-^]_i_ is time-dependent and [Cl^-^]_o_ is constant, to

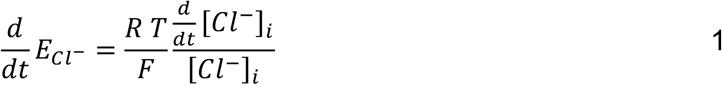

From the current to concentration equation:

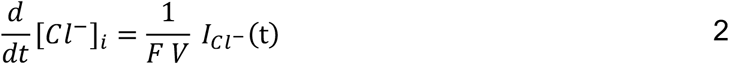

with

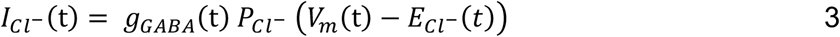

Therefore,

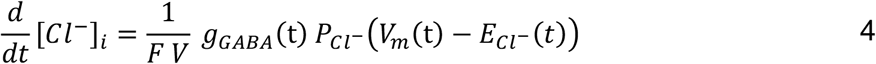

And substituting Eq. 4 into Eq. 1:

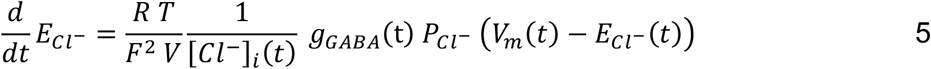

Instead of using [Cl^-^]_i_, we can keep the equation in terms of E_Cl_ by rearranging the Nernst equation for E_Cl_ in terms of [Cl^-^]_i_ (we omit (t) for clarity):

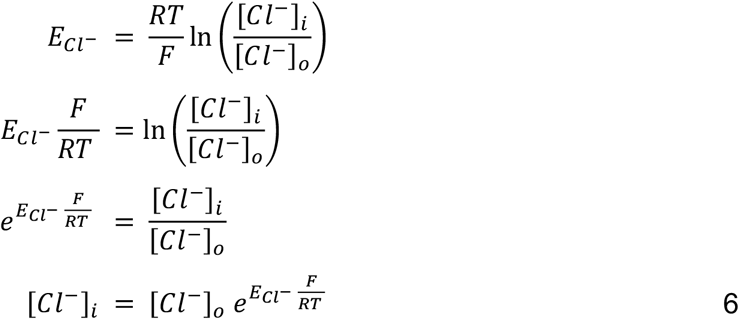

which substituting Eq. 6 into Eq. 5 results in

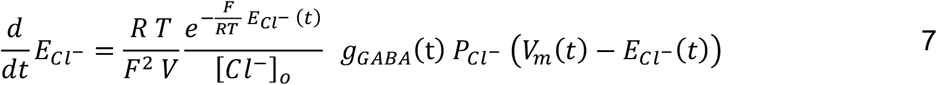

Simplifying by substituting Eq. 3 and 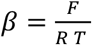, then

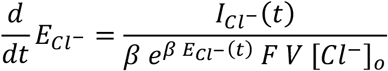

For completeness, we can solve for 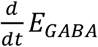 too. Given 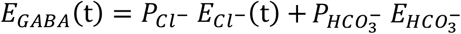, with condition that 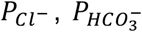, and 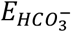 are assumed constant in these simulations, then

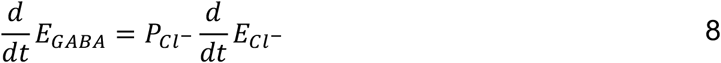

Therefore, by substituting Eq. 7 into Eq. 8 yields

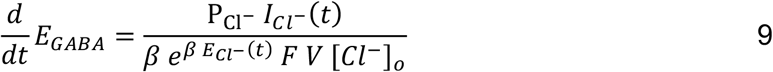

### S2. Activity-based calculation of KCC2’s time constant (τ_KCC2_)

In this work KCC2’s time constant (τ_KCC2_) was set at specific values and the network activity allowed to evolve over the duration of the numerical simulations. Here, I explore a semi-analytical approach for determining what τ_KCC2_ should be, given inhibitory synaptic activity onto a neuron, for keeping ECl close to its hyperpolarising steady-state value.

As a reminder,

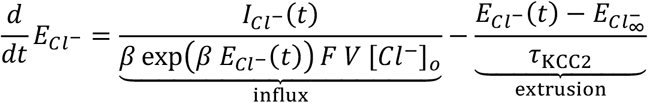

with

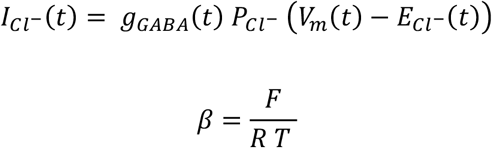

For ease, we do one additional substitution for the influx terms that remain constant, namely

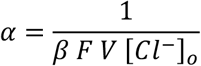

to yield

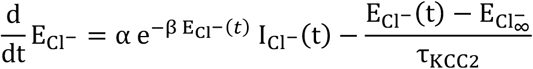

If we want influx and efflux to be balanced (over Δ*t*),

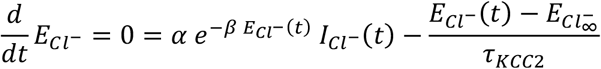

then

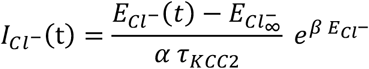

or

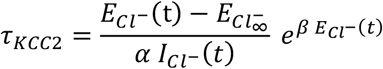

Example:

The time course for a single GABA_A_R’s conductance after receiving a presynaptic action potential is

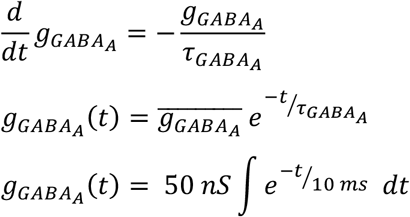

where 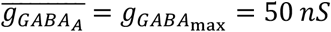 and 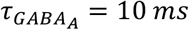.

For 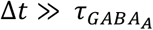,

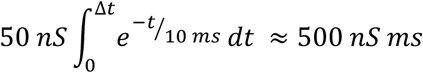

The current, 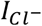, at time *t* is

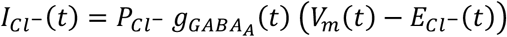

For a single synaptic event lasting Δ*t*, the total current is

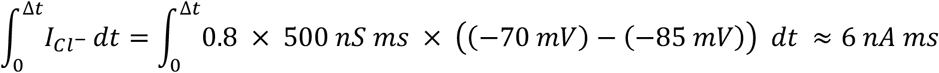

For an input frequency of 10 Hz from 8 inhibitory synapses, the total current is

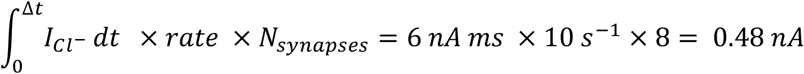

we can now calculate how much change in 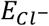 resulted from 0.48 *nA* input, given no extrusion.

With 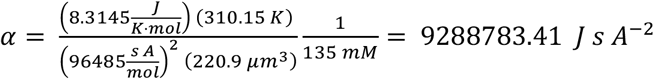

And *β* = 37.42 *J s A*

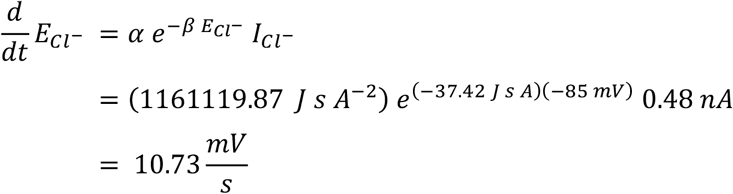

Thus, the new 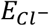 would be -74.27 mV. And the extrusion time constant needed to balance such a change (and return to -85 mV 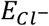) would be

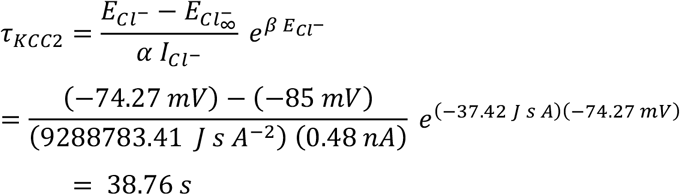

### S3. Drivers of oscillatory network bursts in a spiking neural network model of status epilepticus

*In vitro* models of seizure-like events (SLEs) and status epilepticus (SE) use a 0 mM magnesium concentration solution (0 [Mg^2+^]) to de-inactivate NMDA receptors and enhance excitation in brain slices. This method has historically been used to uncover novel insights and therapeutic results related to seizure control. Experimental evidence has indicated that E_GABA_ can dramatically shift from a baseline of ≈ -85 mV to -40 mV during LRD *(14)*.

To investigate the E_GABA_-dependency of firing rate bursts in the spiking neural network model, under 0 [Mg^2+^] conditions (Figure S1A), E_GABA_ in all cells was incremented by 2 mV every 60 seconds from -74 mV until -42 mV (Figure S1B). The network’s population rate started to exhibit half-second bursts from around -60 to -58 mV (Figure S1B), which are comparable to an experimental model of SE *(14)*. A burst was defined as the average population rate deviating by more than two standard deviations and reaching above 20 Hz. An example of a burst (Figure S1B inset) shows the inter-burst interval (≈ 9.23 s), the burst duration (≈ 0.61 s), as well as the amplitude of the firing rate from baseline for the different neuron populations (PC ≈ 52.44 Hz, red trace; IN ≈ 67.52 Hz, blue trace). The average amplitude for the entire population (dark grey line) was ≈ 56.26 Hz. As E_GABA_ became increasingly depolarised, bursts increased in amplitude (IN up to ≈ 85 Hz, PC up to ≈ 67 Hz) and occurrence (up to every ≈ 9 seconds). Bursts appeared to be driven by large NMDA conductances (Figure S1C, orange trace), with a peak in the population rate burst corresponding with a peak in g_NMDA_ (Figure S1B and C, insets grey vertical line). In contrast, GABA and AMPA peak conductances are before the peak population rate.

A finite vesicle pool, x_s_, prevented runaway activity and halted bursts (Figure S1D). The glutamate vesicle availability 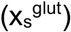, whether targeted onto PC (E → E, Figure S1D, pink trace) or IN (E → I, Figure S1F, red trace) populations facilitated the termination of bursts. In fact, the parabolic shape of the population rate during bursts closely matches the change in 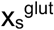 during the same period, with the inflection point of the 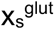 corresponding with the peak of a burst. Furthermore, the GABA vesicle availability 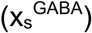, whether onto PC (I → E, Figure S1D, blue trace) or IN (I → I, Figure S1D turquoise trace) neurons, started rapidly dropping before 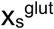 (Figure S1 inset, coloured arrows). Together, the g_NMDA_, x_s_, and synaptic utilisation (traces not shown) parameters determine the duration and interval for bursts, but g_GABA_ and E_GABA_ determine whether they occur.

To assess the effect of network size, the number of neurons was increased while the maximum synaptic conductances were scaled according to 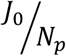 *(43)* where J_0_ = 1000 and N_p_ is the size of the neuron population. Simulations with 1000, 5000, or 10000 neurons showed similar overall neuronal activity (Figure S1E) as well as population burst profiles (Figure S1E inset). Minor differences included lower variability in the population rate, larger amplitude in the bursts, and more depolarised E_GABA_ for bursts to start occurring.

Additional robustness checks were performed on the external input and leak conductances such that similar network activity was observed for g_leak_ = 1 nS, g_AMPAmax,2Hz_ = 1 nS, and only 100 external Poisson inputs (Figure S1F). For this alternative parameter set, network bursts started at a less depolarised E_GABA_ of -66 mV (instead of -60 for the default parameter set) and had a higher burst amplitude (Figure S1F, inset).

Finally, the maximum conductances for AMPA and NMDA were also explored to assess their influence on network bursting behaviour (Figure S2). As this work has rigorously explored the paradoxical effect of E_GABA_ and g_GABAmax_ on network bursting, g_AMPAmax_ and g_NMDAmax_ were assessed at three values of E_GABA_ (−42, -56, and -70 mV) and three values of g_GABAmax_ (25, 50, and 100 nS). We found that increasing g_AMPAmax_ had a relatively weak effect on the number of bursts (compared to g_GABAmax_ for example), with its biggest influence occurring at small number of bursts (e.g. first columns in Figure S2B and C); the increased excitation pushes a network on the brink of a synchronised event to actually occur. Unsurprisingly due to its long time constant, g_NMDAmax_ had a substantial impact on the number of bursts in the network at all E_GABA_ and g_GABAmax_ levels. Increasing g_NMDAmax_ could cause the network to burst at least an additional 2 times and up to 8 more times at low E_GABA_ (Figure S2C, first column). The selected maximum NMDA conductance (g_NMDAmax_) of 5 nS proved appropriate for inducing bursts at specific GABA_A_ reversal potentials (E_GABA_) and maximum GABA conductances (g_GABAmax_), while still allowing for a balanced influence on network dynamics. This choice of g_NMDAmax_ significantly affected population firing rates without overshadowing other key network behaviors.

## ACKNOWLEDGEMENTS

The research leading to these results has received support from the National Research Foundation of South Africa, the German Deutscher Akademischer Austauschdienst, NOMIS Foundation, NVIDIA Academic Program, the University of Cape Town, the Anna Mueller Grocholski Foundation, the Swiss National Science Foundation (SNSF: 208184), the Gabriel Foundation, a Wellcome Trust Seed Award (214042/Z/18/Z), the South African Medical Research Council and by the FLAIR Fellowship Programme (FLR\R1\190829): a partnership between the African Academy of Sciences and the Royal Society funded by the UK Government’s Global Challenges Research Fund and a Wellcome Trust International Intermediate Fellowship (222968/Z/21/Z).

## CONTRIBUTIONS

### Author contributions

Conceptualization: CBC, JVR

Methodology: CBC, HS

Investigation: CBC, RJB, TF, GR, RER

Visualization: RJB, CBC

Funding acquisition: JVR, CBC

Project administration: JVR

Supervision: JVR, HS

Writing – original draft: CBC, RJB

Writing – review & editing: All authors.

### Data and materials availability

All data are available in the main text or the supplementary materials.

## Supplementary figures

**Figure S1:**
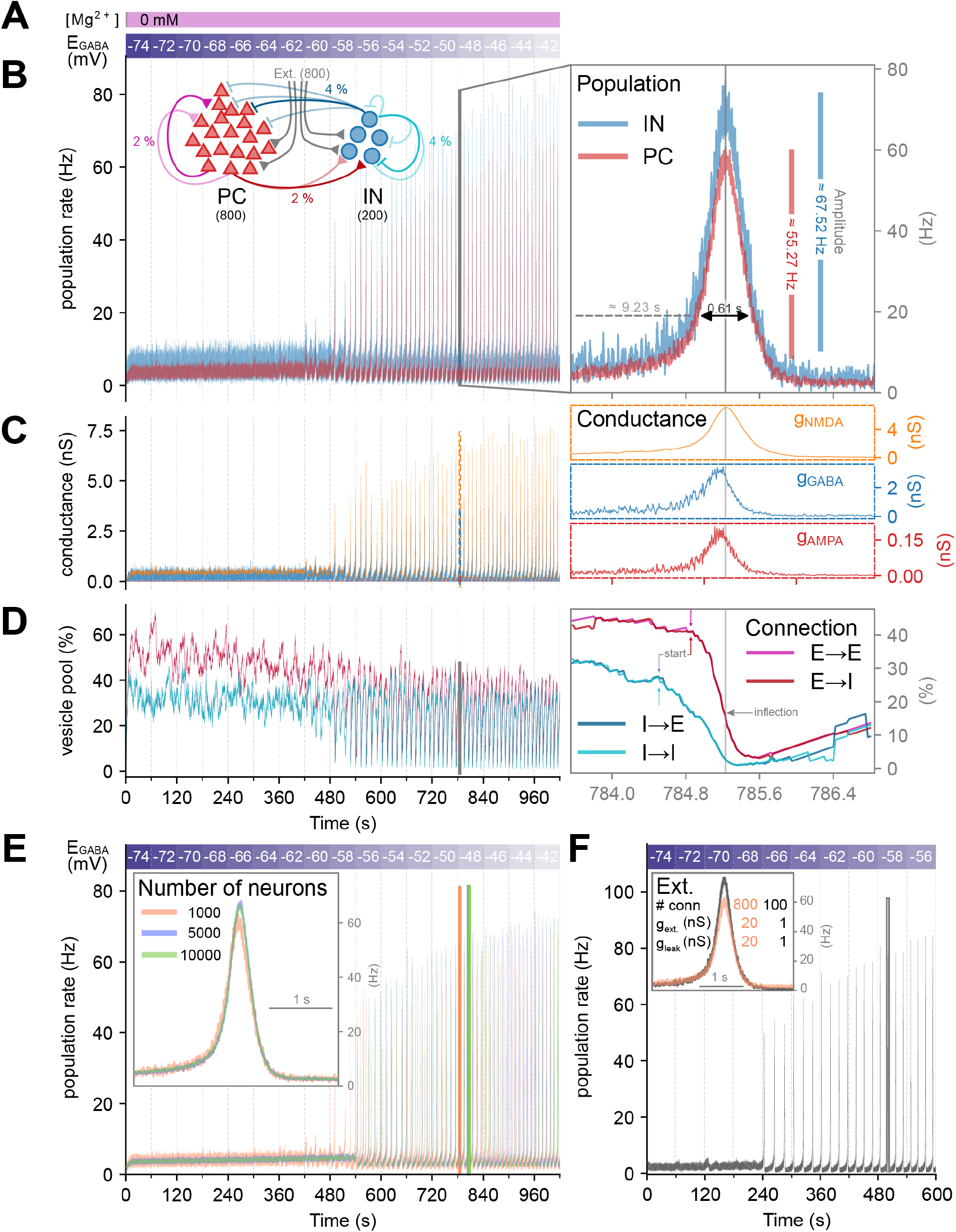
A spiking neural network model of status epilepticus. **(A)** The Mg^2+^ block typically associated with NMDA inactivation was removed (“de-inactivated”) by changing the concentration of Mg^2+^ to 0 as done experimentally *(14)*. E_GABA_ was varied at discrete time points, every 60 seconds, by 2 mV, from -74 mV until -42 mV. **(B, inset)** Neuron populations and associated connections. The network consisted of 800 pyramidal cells (PC, red), 200 interneurons (IN, blue). Each PC had a 2 % probability of synapsing onto another PC (p_EE_ = 0.02, E → E, brown) or onto an interneuron (p_IE_ = 0.02, E → I, yellow). Each IN had a 4% probability of synapsing onto another IN (p_II_ = 0.04, I → I, turquoise) or onto a PC (p_EI_ = 0.04, I → E, purple). **(B)** Mean instantaneous firing rate (time window = 10.1 ms) for PC (red) and IN (blue) populations. Right, an example of a burst, which typically latest 0.61 s and occurred 9.23 s after the previous burst ended. The burst amplitudes for the IN (blue, 67.52 Hz), and PC (red, 55.27 Hz) populations are shown. The IN has the highest mean firing rate and reaches the highest peak firing rate. **(C)** Mean conductances for 4 neurons, separated into NMDA (orange), AMPA (red), and GABA (type A, blue) receptor conductances. Right, individual conductances during a burst. Parameters for max conductances were as follows: g_AMPAmax_ = 5 nS, g_NMDAmax_ = 5 nS, g_GABAmax_ = 50 nS. **(D)** Examples of vesicle pools (x_s_) for 4 neurons, separated into E → E (pink), E → I (red), I → I (turquoise), and I → E (blue) connections. Right, the depletion of vesicles during an oscillation. Arrows indicate the start of rapid vesicle depletion during the time window or the point of inflection (grey) for the glutamatergic vesicle pool (E → E and E → I). **(E)** Networks of 5000 (blue) and 10000 (green) neurons (of which, 20% IN) produced similar results to a network with 1000 (orange) neurons when the max synaptic conductances were multiplied by 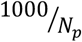 where N_p_ is the number of neurons in the population. Networks with more neurons had less population rate variability but similar mean, minimum, and maximum values. Inset, highlight of an example burst for each population at E_GABA_ = -48 mV. **(F)** Decreasing the number (# conn) and strength (g_ext_) of the external connections (Ext.), along with a weaker leak conductance (g_leak_) also produced comparable results, albeit with network bursting starting at a less depolarised E_GABA_ and larger amplitude network brusts. Inset, network bursts of the default parameter set used throughout the work (orange, E_GABA_ = - 48 mV) and the alternative set of parameters (grey, E_GABA_ = - 58 mV).

**Figure S2:**
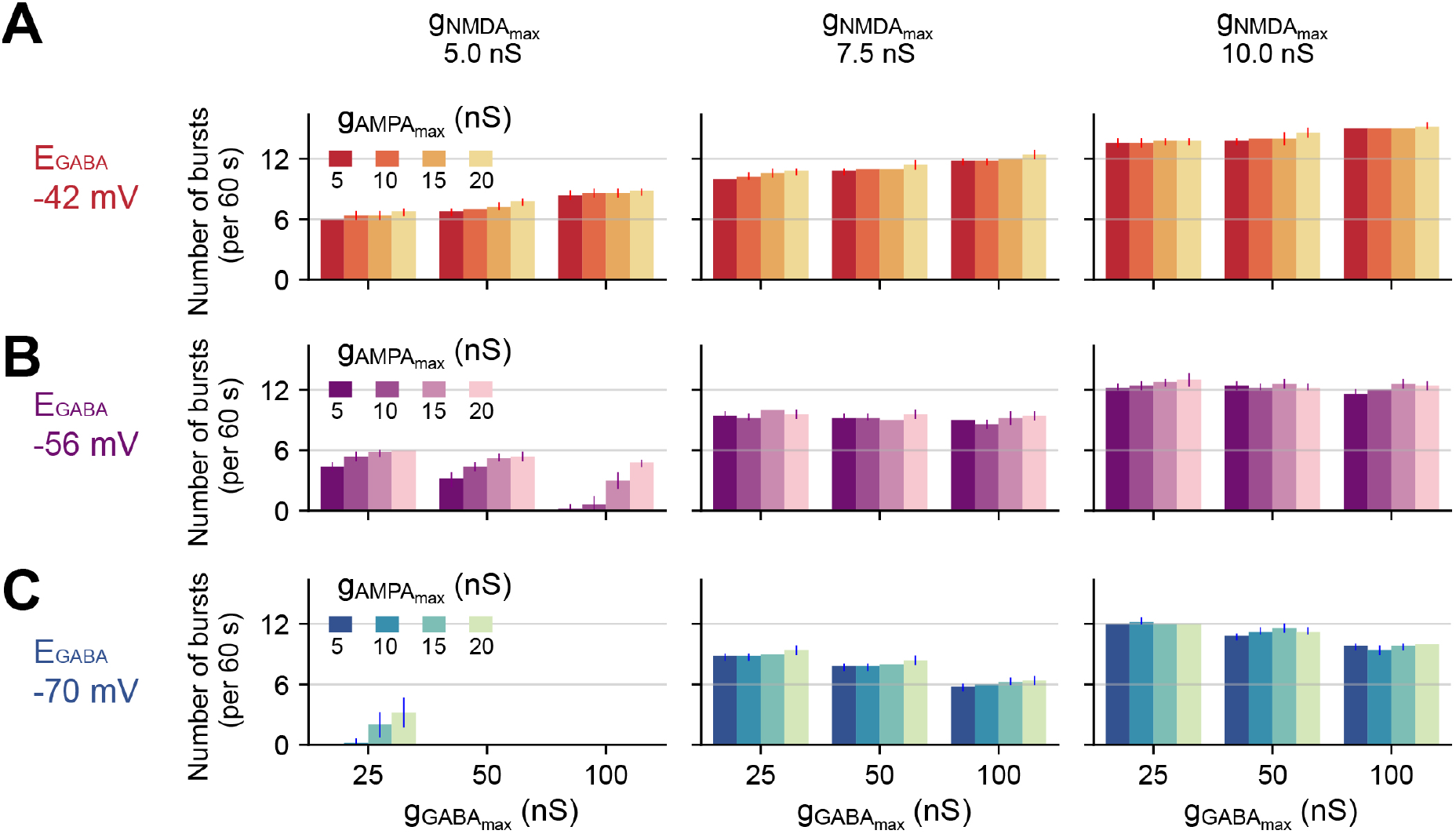
The additional influence of NMDA and AMPA conductances on the number of bursts. Per plot, the maximum GABA conductance was varied on the x-axis to be 25, 50, or 100 nS and the number of bursts (per 60 s) recorded. In addition, the maximum AMPA conductance (g_AMPAmax_) was set to either 5, 10, 15, or 20 nS (brightness). Per column, the maximum NMDA conductance (g_NMDAmax_) was set to either 5, 7.5, or 10 nS. Per row, the GABA_A_R reversal potential (E_GABA_) was set to -70 **(C)**, -56 **(B)**, or -42 **(A)** mV.

## REFERENCES

1. E. Trinka, H. Cock, D. Hesdorffer, A. O. Rossetti, I. E. Scheffer, S. Shinnar, S. Shorvon, D. H. Lowenstein, A definition and classification of status epilepticus - Report of the ILAE Task Force on Classification of Status Epilepticus. Epilepsia 56, 1515–1523 (2015).

2. J. G. Boggs, Mortality associated with status epilepticus. Epilepsy Currents 4, 25–27 (2004).

3. T. Glauser, S. Shinnar, D. Gloss, B. Alldredge, R. Arya, J. Bainbridge, M. Bare, T. Bleck, W. Edwin Dodson, L. Garrity, A. Jagoda, D. Lowenstein, J. Pellock, J. Riviello, E. Sloan, D. M. Treiman, Evidence-based guideline: Treatment of convulsive status epilepticus in children and adults: Report of the guideline committee of the American epilepsy society. Epilepsy Currents 16, 48–61 (2016).

4. R. Appleton, I. Choonara, T. Martland, B. Phillips, R. Scott, W. Whitehouse, The treatment of convulsive status epilepticus in children. Archives of Disease in Childhood 83, 415–419 (2000).

5. S. A. Mayer, J. Claassen, J. Lokin, F. Mendelsohn, L. J. Dennis, B. F. Fitzsimmons, Refractory status epilepticus: Frequency, risk factors, and impact on outcome. Archives of Neurology 59, 205–210 (2002).

6. R. F. Chin, B. G. Neville, C. Peckham, A. Wade, H. Bedford, R. C. Scott, Treatment of community-onset, childhood convulsive status epilepticus: a prospective, population-based study. The Lancet Neurology 7, 696–703 (2008).

7. R. J. Burman, R. E. Rosch, J. M. Wilmshurst, A. Sen, G. Ramantani, C. J. Akerman, J. V. Raimondo, Why won’t it stop? The dynamics of benzodiazepine resistance in status epilepticus. Nature Reviews Neurology (2022), doi:10.1038/s41582-022-00664-3.

8. A. J. Trevelyan, C. a Schevon, How inhibition influences seizure propagation. Neuropharmacology 69, 45–54 (2013).

9. J. V. Raimondo, B. A. Richards, M. A. Woodin, Neuronal chloride and excitability — the big impact of small changes. Current Opinion in Neurobiology 43, 35–42 (2017).

10. A. G. Chapman, B. S. Meldrum, B. K. Siesiö, Cerebral Metabolic Changes During Prolonged Epileptic Seizures in Rats. Journal of Neurochemistry 28, 1025–1035 (1977).

11. A. Pereira de Vasconcelos, G. el Hamdi, P. Vert, A. Nehlig, An experimental model of generalized seizures for the measurement of local cerebral glucose utilization in the immature rat. II. Mapping of brain metabolism using the quantitative [14C]2-deoxyglucose technique. Brain Res Dev Brain Res 69, 243–259 (1992).

12. A. Nehlig, A. Pereira de Vasconcelos, The model of pentylenetetrazol-induced status epilepticus in the immature rat: short- and long-term effects. Epilepsy Res 26, 93–103 (1996).

13. A. Pitkanen, P. Schwartzkroin, S. Moshe, Models of Seizures and Epilepsy (2006; https://books.google.co.za/books?hl=en&lr=&id=qUXUDQAAQBAJ&oi=fnd&pg=PP1&dq=Models+of+Seizures+and+Epilepsy&ots=t56bP35Mwq&sig=_ooevDMUzoMOAdmuztwLNEEbp3k#v=onepage&q=ModelsofSeizuresandEpilepsy&f=false).

14. R. J. Burman, J. S. Selfe, J. H. Lee, M. van den Berg, A. Calin, N. K. Codadu, R. Wright, S. E. Newey, R. R. Parrish, A. A. Katz, J. M. Wilmshurst, C. J. Akerman, A. J. Trevelyan, J. V. Raimondo, Excitatory GABAergic signalling is associated with benzodiazepine resistance in status epilepticus. Brain 142, 3482–3501 (2019).

15. G. El Hamdi, A. Pereira de Vasconcelos, P. Vert, A. Nehlig, An experimental model of generalized seizures for the measurement of local cerebral glucose utilization in the immature rat. I. Behavioral characterization and determination of lumped constant. Developmental Brain Research 69, 233–242 (1992).

16. C. J. Rogers, R. E. Twyman, R. L. Macdonald, Benzodiazepine and beta-carboline regulation of single GABAA receptor channels of mouse spinal neurones in culture. J Physiol 475, 69–82 (1994).

17. T. R. Browne, J. K. Penry, Benzodiazepines in the Treatment of Epilepsy A Review. Epilepsia 14, 277–310 (1973).

18. M. Farrant, K. Kaila, The cellular, molecular and ionic basis of GABAA receptor signalling. Progress in Brain Research 160, 59–87 (2007).

19. J. V. Raimondo, H. Markram, C. J. Akerman, Short-term ionic plasticity at GABAergic synapses. Front. Synaptic Neurosci. 4, 1–9 (2012).

20. K. Kaila, T. J. Price, J. a. Payne, M. Puskarjov, J. Voipio, Cation-chloride cotransporters in neuronal development, plasticity and disease. Nature Reviews Neuroscience 15, 637–654 (2014).

21. Y. Ben-Ari, Excitatory actions of GABA during development: the nature of the nurture. Nat. Rev. Neurosci. 3, 728–739 (2002).

22. G. Huberfeld, L. Wittner, S. Clemenceau, M. Baulac, K. Kaila, R. Miles, C. Rivera, Perturbed chloride homeostasis and GABAergic signaling in human temporal lobe epilepsy. J. Neurosci. 27, 9866–73 (2007).

23. T. J. Ellender, J. V. Raimondo, A. Irkle, K. P. Lamsa, C. J. Akerman, Excitatory effects of parvalbumin-expressing interneurons maintain hippocampal epileptiform activity via synchronous afterdischarges. The Journal of neuroscience : the official journal of the Society for Neuroscience 34, 15208–22 (2014).

24. S. Sulis Sato, P. Artoni, S. Landi, O. Cozzolino, R. Parra, E. Pracucci, F. Trovato, J. Szczurkowska, S. Luin, D. Arosio, F. Beltram, L. Cancedda, K. Kaila, G. M. Ratto, Simultaneous two-photon imaging of intracellular chloride concentration and pH in mouse pyramidal neurons in vivo. Proceedings of the National Academy of Sciences, 201702861 (2017).

25. V. Magloire, J. Cornford, A. Lieb, D. M. Kullmann, I. Pavlov, KCC2 overexpression prevents the paradoxical seizure-promoting action of somatic inhibition. Nature Communications 10, 1225 (2019).

26. R. Miles, R. K. S. Wong, R. D. Traub, Synchronized afterdischarges in the hippocampus: Contribution of local synaptic interactions. Neuroscience 12, 1179–1189 (1984).

27. T. Viitanen, E. Ruusuvuori, K. Kaila, J. Voipio, The K+-Cl-cotransporter KCC2 promotes GABAergic excitation in the mature rat hippocampus. The Journal of Physiology 588, 1527–1540 (2010).

28. M. Wenzel, J. P. Hamm, D. S. Peterka, R. Y. Correspondence, R. Yuste, Reliable and Elastic Propagation of Cortical Seizures In Vivo. Cell Reports 19, 2681–2693 (2017).

29. S. Sivakumaran, J. Maguire, Bumetanide reduces seizure progression and the development of pharmacoresistant status epilepticus. Epilepsia 57, 222–232 (2016).

30. D. L. Cheung, M. J. Cooke, C. S. Goulton, C. Chaichim, L. F. Cheung, A. Khoshaba, J. Nabekura, A. J. Moorhouse, Global transgenic upregulation of KCC2 confers enhanced diazepam efficacy in treating sustained seizures. Epilepsia 63, e15–e22 (2022).

31. R. Jarvis, S. F. Josephine Ng, A. J. Nathanson, R. A. Cardarelli, K. Abiraman, F. Wade, A. Evans-Strong, M. P. Fernandez-Campa, T. Z. Deeb, J. L. Smalley, T. Jamier, I. K. Gurrell, L. McWilliams, A. Kawatkar, L. C. Conway, Q. Wang, R. W. Burli, N. J. Brandon, I. P. Chessell, A. J. Goldman, J. L. Maguire, S. J. Moss, Direct activation of KCC2 arrests benzodiazepine refractory status epilepticus and limits the subsequent neuronal injury in mice. Cell Reports Medicine 4, 100957 (2023).

32. N. Doyon, S. a Prescott, A. Castonguay, A. G. Godin, H. Kröger, Y. De Koninck, Efficacy of synaptic inhibition depends on multiple, dynamically interacting mechanisms implicated in chloride homeostasis. PLoS computational biology 7, 1–22 (2011).

33. P. Jedlicka, T. Deller, B. Gutkin, Activity dependent intracellular chloride accumulation and diffusion controls GABAA receptor mediated synaptic transmission. Hippocampus 898, 885–898 (2011).

34. C. B. Currin, A. J. Trevelyan, C. J. Akerman, J. V. Raimondo, H. Berry, Ed. Chloride dynamics alter the input-output properties of neurons. PLoS Computational Biology 16, e1007932 (2020).

35. C. B. Currin, J. V. Raimondo, J. Rubin, Ed. Computational models reveal how chloride dynamics determine the optimal distribution of inhibitory synapses to minimise dendritic excitability. PLoS Comput Biol 18, e1010534 (2022).

36. T. Fedele, R. J. Burman, A. Steinberg, G. Selmin, G. Ramantani, R. E. Rosch, Synaptic inhibitory dynamics drive benzodiazepine response in paediatric status epilepticus, 2023.08.23.23294456 (2023).

37. K. Staley, Enhancement of the excitatory actions of GABA by barbiturates and benzodiazepines. Neuroscience Letters 146, 105–107 (1992).

38. T. Z. Deeb, J. Maguire, S. J. Moss, Possible alterations in GABAA receptor signaling that underlie benzodiazepine-resistant seizures. Epilepsia 53, 79–88 (2012).

39. L. S. Deshpande, R. E. Blair, N. Nagarkatti, S. Sombati, B. R. Martin, R. J. DeLorenzo, Development of pharmacoresistance to benzodiazepines but not cannabinoids in the hippocampal neuronal culture model of status epilepticus. Exp Neurol 204, 705–713 (2007).

40. N. K. Codadu, R. T. Graham, R. J. Burman, R. T. Jackson-Taylor, J. V. Raimondo, A. J. Trevelyan, R. R. Parrish, Divergent paths to seizure-like events. Physiological Reports 7, 1–15 (2019).

41. V. Magloire, J. Cornford, A. Lieb, D. M. Kullmann, I. Pavlov, KCC2 overexpression prevents the paradoxical seizure-promoting action of somatic inhibition. Nature Communications 10, 1–13 (2019).

42. O. Melamed, O. Barak, G. Silberberg, H. Markram, M. Tsodyks, Slow oscillations in neural networks with facilitating synapses. Journal of Computational Neuroscience 25, 308–316 (2008).

43. W. Gerstner, W. M. Kistler, R. Naud, L. Paninski, Neuronal Dynamics W. Gerstner, W. M. Kistler, R. Naud, L. Paninski, Eds. (Cambridge University Press, Cambridge, 2014; https://neuronaldynamics.epfl.ch/index.html).

44. R. J. Burman, J. V. Raimondo, J. G. R. Jefferys, A. Sen, C. J. Akerman, The transition to status epilepticus: how the brain meets the demands of perpetual seizure activity. Seizure 75, 137–144 (2020).

45. Y. Wang, Y. Wang, Z. Chen, Double-edged GABAergic synaptic transmission in seizures: The importance of chloride plasticity. Brain Research 1701, 126–136 (2018).

46. C. Rivera, J. Voipio, J. Thomas-Crusells, H. Li, Z. Emri, S. Sipilä, J. A. Payne, L. Minichiello, M. Saarma, K. Kaila, Mechanism of activity-dependent downregulation of the neuron-specific K-Cl cotransporter KCC2. J Neurosci. 24, 4683–91 (2004).

47. H. H. C. Lee, T. Z. Deeb, J. A. Walker, P. A. Davies, S. J. Moss, NMDA receptor activity downregulates KCC2 resulting in depolarizing GABAA receptor–mediated currents. Nature Neuroscience 14, 736–743 (2011).

48. M. Gaínza-Lein, I. Sánchez Fernández, M. Jackson, N. S. Abend, R. Arya, J. N. Brenton, J. L. Carpenter, K. E. Chapman, W. D. Gaillard, T. A. Glauser, J. L. Goldstein, H. P. Goodkin, K. Kapur, M. A. Mikati, K. Peariso, R. C. Tasker, D. Tchapyjnikov, A. A. Topjian, M. S. Wainwright, A. Wilfong, K. Williams, T. Loddenkemper, Pediatric Status Epilepticus Research Group, Association of Time to Treatment With Short-term Outcomes for Pediatric Patients With Refractory Convulsive Status Epilepticus. JAMA Neurol 75, 410–418 (2018).

49. J. Kapur, D. a Coulter, Experimental status epilepticus alters gamma-aminobutyric acid type A receptor function in CA1 pyramidal neurons. Annals of neurology 38, 893–900 (1995).

50. H. P. Goodkin, J.-L. Yeh, J. Kapur, Status epilepticus increases the intracellular accumulation of GABAA receptors. The Journal of neuroscience : the official journal of the Society for Neuroscience 25, 5511–20 (2005).

51. H. P. Goodkin, S. Joshi, Z. Mtchedlishvili, J. Brar, J. Kapur, Subunit-Specific Trafficking of GABAA Receptors during Status Epilepticus. Journal of Neuroscience 28, 2527–2538 (2008).

52. H. Alfonsa, J. H. Lakey, R. N. Lightowlers, A. J. Trevelyan, Cl-out is a novel cooperative optogenetic tool for extruding chloride from neurons. Nature communications 7 (2016), doi:10.1038/ncomms13495.

53. E. McMoneagle, J. Zhou, S. Zhang, W. Huang, S. S. Josiah, K. Ding, Y. Wang, J. Zhang, Neuronal K+-Cl-cotransporter KCC2 as a promising drug target for epilepsy treatment. Acta Pharmacol Sin 45, 1–22 (2024).

54. S. Sivakumaran, R. A. Cardarelli, J. Maguire, M. R. Kelley, L. Silayeva, D. H. Morrow, J. Mukherjee, Y. E. Moore, R. J. Mather, M. E. Duggan, N. J. Brandon, J. Dunlop, S. Zicha, S. J. Moss, T. Z. Deeb, Selective inhibition of KCC2 leads to hyperexcitability and epileptiform discharges in hippocampal slices and in vivo. Journal of Neuroscience 35, 8291–8296 (2015).

55. M. A. Kramer, W. Truccolo, U. T. Eden, K. Q. Lepage, L. R. Hochberg, E. N. Eskandar, Joseph. R. Madsen, J. W. Lee, A. Maheshwari, E. Halgren, C. J. Chu, S. S. Cash, Human seizures self-terminate across spatial scales via a critical transition. Proceedings of the National Academy of Sciences 109, 21116–21121 (2012).

56. G. P. Krishnan, M. Bazhenov, Ionic dynamics mediate spontaneous termination of seizures and postictal depression state. The Journal of neuroscience : the official journal of the Society for Neuroscience 31, 8870–82 (2011).

57. F. Fröhlich, M. Bazhenov, I. Timofeev, T. J. Sejnowski, Maintenance and termination of neocortical oscillations by dynamic modulation of intrinsic and synaptic excitability. Thalamus & related systems 3, 147–156 (2005).

58. J. V. Raimondo, R. J. Burman, A. A. Katz, C. J. Akerman, Ion dynamics during seizures. Frontiers in Cellular Neuroscience 9, 1–14 (2015).

59. R. L. Macdonald, J. L. Barker, Different actions of anticonvulsant and anesthetic barbiturates revealed by use of cultured mammalian neurons. Science 200, 775–777 (1978).

60. R. Nardou, S. Yamamoto, A. Bhar, N. Burnashev, Y. Ben-Ari, I. Khalilov, Phenobarbital but Not Diazepam Reduces AMPA/kainate Receptor Mediated Currents and Exerts Opposite Actions on Initial Seizures in the Neonatal Rat Hippocampus. Frontiers in Cellular Neuroscience 5, 1–16 (2011).

61. G. R. Lee, R. Gommers, F. Waselewski, K. Wohlfahrt, A. O’Leary, PyWavelets: A Python package for wavelet analysis. Journal of Open Source Software 4, 1237 (2019).

62. W. W. Anderson, D. V. Lewis, H. S. Swartzwelder, W. A. Wilson, Magnesium-free medium activates seizure-like events in the rat hippocampal slice. Brain Res 398, 215–219 (1986).

63. J. P. Dreier, U. Heinemann, Regional and time dependent variations of low Mg2+ induced epileptiform activity in rat temporal cortex slices. Exp Brain Res 87, 581–596 (1991).

64. T. J. Ellender, J. V. Raimondo, A. Irkle, K. P. Lamsa, C. J. Akerman, Excitatory effects of parvalbumin-expressing interneurons maintain hippocampal epileptiform activity via synchronous afterdischarges. Journal of Neuroscience 34, 15208–15222 (2014).

65. N. Brunel, Phase diagrams of sparsely connected networks of excitatory and inhibitory spiking neurons. Neurocomputing 32–33, 307–312 (2000).

66. T. P. Vogels, H. Sprekeler, F. Zenke, C. Clopath, W. Gerstner, Inhibitory plasticity balances excitation and inhibition in sensory pathways and memory networks. Science 334, 1569–73 (2011).

67. N. Parga, L. F. Abbott, Network model of spontaneous activity exhibiting synchronous transitions between up and down states. Frontiers in Neuroscience 1, 57–66 (2007).

68. J. A. Wolf, J. T. Moyer, M. T. Lazarewicz, D. Contreras, M. Benoit-Marand, P. O’Donnell, L. H. Finkel, NMDA/AMPA ratio impacts state transitions and entrainment to oscillations in a computational model of the nucleus accumbens medium spiny projection neuron. Journal of Neuroscience 25, 9080–9095 (2005).

69. H. Sanders, M. Berends, G. Major, M. S. Goldman, J. E. Lisman, NMDA and GABAB (KIR) conductances: The “perfect couple” for bistability. Journal of Neuroscience 33, 424–429 (2013).

70. I. Timofeev, F. Grenier, M. Bazhenov, T. J. Sejnowski, M. Steriade, Origin of slow cortical oscillations in deafferented cortical slabs. Cerebral Cortex 10, 1185–1199 (2000).

71. M. Tsodyks, H. Markram, The neural code between neocortical pyramidal neurons depends. Proceedings of the National Academy of Sciences of the United States of America 94, 719–723 (1997).

72. P. Jedlicka, T. Deller, B. S. Gutkin, K. H. Backus, Activity-dependent intracellular chloride accumulation and diffusion controls GABA(A) receptor-mediated synaptic transmission. Hippocampus 21, 885–98 (2011).

73. M. Stimberg, R. Brette, D. F. M. Goodman, Brian 2, an intuitive and efficient neural simulator. eLife 8 (2019), doi:10.7554/eLife.47314.

74. C. R. Harris, K. J. Millman, S. J. van der Walt, R. Gommers, P. Virtanen, D. Cournapeau, E. Wieser, J. Taylor, S. Berg, N. J. Smith, R. Kern, M. Picus, S. Hoyer, M. H. van Kerkwijk, M. Brett, A. Haldane, J. F. del Río, M. Wiebe, P. Peterson, P. Gérard-Marchant, K. Sheppard, T. Reddy, W. Weckesser, H. Abbasi, C. Gohlke, T. E. Oliphant, Array programming with NumPy. Nature 585, 357–362 (2020).

75. P. Virtanen, R. Gommers, T. E. Oliphant, M. Haberland, T. Reddy, D. Cournapeau, E. Burovski, P. Peterson, W. Weckesser, J. Bright, S. J. van der Walt, M. Brett, J. Wilson, K. J. Millman, N. Mayorov, A. R. J. Nelson, E. Jones, R. Kern, E. Larson, C. J. Carey, I. Polat, Y. Feng, E. W. Moore, J. VanderPlas, D. Laxalde, J. Perktold, R. Cimrman, I. Henriksen, E. A. Quintero, C. R. Harris, A. M. Archibald, A. H. Ribeiro, F. Pedregosa, P. van Mulbregt, A. Vijaykumar, A. P. Bardelli, A. Rothberg, A. Hilboll, A. Kloeckner, A. Scopatz, A. Lee, A. Rokem, C. N. Woods, C. Fulton, C. Masson, C. Häggström, C. Fitzgerald, D. A. Nicholson, D. R. Hagen, D. V. Pasechnik, E. Olivetti, E. Martin, E. Wieser, F. Silva, F. Lenders, F. Wilhelm, G. Young, G. A. Price, G. L. Ingold, G. E. Allen, G. R. Lee, H. Audren, I. Probst, J. P. Dietrich, J. Silterra, J. T. Webber, J. Slavič, J. Nothman, J. Buchner, J. Kulick, J. L. Schönberger, J. V. de Miranda Cardoso, J. Reimer, J. Harrington, J. L. C. Rodríguez, J. Nunez-Iglesias, J. Kuczynski, K. Tritz, M. Thoma, M. Newville, M. Kümmerer, M. Bolingbroke, M. Tartre, M. Pak, N. J. Smith, N. Nowaczyk, N. Shebanov, O. Pavlyk, P. A. Brodtkorb, P. Lee, R. T. McGibbon, R. Feldbauer, S. Lewis, S. Tygier, S. Sievert, S. Vigna, S. Peterson, S. More, T. Pudlik, T. Oshima, T. J. Pingel, T. P. Robitaille, T. Spura, T. R. Jones, T. Cera, T. Leslie, T. Zito, T. Krauss, U. Upadhyay, Y. O. Halchenko, Y. Vázquez-Baeza, SciPy 1.0: fundamental algorithms for scientific computing in Python. Nature Methods 17, 261–272 (2020).

76. W. McKinney, Data Structures for Statistical Computing in Python. Proceedings of the 9th Python in Science Conference 1697900, 51–56 (2010).

77. J. D. Hunter, Matplotlib: A 2D graphics environment. Computing in Science and Engineering 9, 90–95 (2007).

78. M. Waskom, the seaborn development team, mwaskom/seaborn (2020), doi:10.5281/zenodo.592845.

79. S. Ramaswamy, S. L. Hill, J. G. King, F. Schürmann, Y. Wang, H. Markram, Intrinsic morphological diversity of thick-tufted layer 5 pyramidal neurons ensures robust and invariant properties of in silico synaptic connections. The Journal of Physiology 590, 737–752 (2012).

80. P. Rhodes, The properties and implications of NMDA spikes in neocortical pyramidal cells. Journal of Neuroscience 26, 6704–6715 (2006).

